# PyEuk: a tool suite for catalogue-free multilocus typing

**DOI:** 10.64898/2026.09.10.750732

**Authors:** Sergei Kosakovsky Pond, Danielle Callan, Anton Nekrutenko

## Abstract

Multilocus sequence typing anchors molecular epidemiology, but traditional frameworks require centrally curated allele catalogues. For emerging and uncultivable eukaryotic parasites, maintaining these databases is impractical, leaving surveillance reliant on fragmented, assay-specific scripts. PyEuk eliminates this bottleneck by providing an open, catalogue-free suite that calls microhaplotypes directly from sequence differences relative to a reference within data-defined genomic windows. When amplicon coordinates are uncharacterized or unpublished, PyEuk reconstructs target panels *de novo* from raw read coverage peaks mapped to a draft assembly. Across benchmark cohorts spanning *Cyclospora cayetanensis* and *Plasmodium vivax*, PyEuk recovers epidemiological structure established by tracebacks, geography, and clinical recurrence without organism-specific tuning. In foodborne outbreaks, it resolves independent transmission chains using either curated or *de novo* panels and scales to national surveillance archives exceeding 8,058 isolates. In *P. vivax* malaria, its heterozygosity-weighted identity-by-state (wIBS) distance separates continental lineages, discriminates liver-stage relapses from reinfections, and delineates transmission clusters. Rather than forcing an arbitrary partition on continuous variation, PyEuk evaluates bootstrap stability, reporting supported cluster count ranges alongside reproducible transmission cores. PyEuk provides a portable, reproducible foundation for eukaryotic pathogen surveillance.

## 1 Introduction

Multilocus sequence typing (MLST) has anchored molecular epidemiology and public health surveillance since 1998. Developed by Maiden et al. (1998) to overcome the inter-laboratory variability and subjective scoring of gel electrophoresis, MLST replaced analog band patterns with digital sequence types. By sequencing internal fragments of conserved housekeeping loci and assigning sequential integers to distinct alleles, the method established an objective, portable classification framework that underpins surveillance and outbreak investigations across hundreds of bacterial, fungal, and parasitic species (Maiden 2006, Jolley et al. 2018).

Even in the era of high-throughput sequencing, MLST and its gene-by-gene descendants remain central. While whole-genome sequencing (WGS) offers complete genome resolution, public health surveillance routinely confronts clinical and environmental specimens—such as stool or produce washes—where pathogen biomass is dwarfed by host and commensal DNA. Under these conditions, unbiased WGS is cost-prohibitive without culture enrichment, whereas targeted amplicon sequencing delivers deep coverage directly from uncultured, low-biomass templates (Neafsey et al. 2021). Furthermore, many clinically significant infections are polyclonal; while short-read WGS collapses co-circulating lineages into chimeric consensus sequences, amplicon deep sequencing recovers physically phased microhaplotypes at single-molecule resolution (Callahan et al. 2016, Callahan et al. 2017, Hathaway et al. 2018, Tessema et al. 2022). High-throughput sequencing accelerated gene-by-gene typing, scaling the approach to core-genome (cgMLST) and whole-genome (wgMLST) schemes spanning thousands of loci (Maiden et al. 2013, Moura et al. 2016, Silva et al. 2018). However, this portability requires centralized curation and database synchronization for every new allele and profile (Uelze et al. 2020). For eukaryotic parasites lacking dedicated consortia, this dependency creates an operational bottleneck.

The foodborne apicomplexan parasite *Cyclospora cayetanensis* illustrates the practical limits of catalogue-dependent typing. Outbreak investigations are bedeviled by prolonged incubation periods and perishable produce vehicles that disperse cases across jurisdictions (Herwaldt 2000, Markon et al. 2025), leaving an orphaned backlog of over 2,200 U.S. cases unlinked between 2011 and 2015 without a validated typing tool (Casillas et al. 2019). Initial five-locus microsatellite typing resolved only 17 genotypes across 54 patients, suffered amplification dropouts, and conflated unrelated cases (Hofstetter et al. 2019). To improve resolution, the Centers for Disease Control and Prevention (CDC) developed a targeted amplicon panel of polymorphic nuclear markers (Houghton et al. 2020). Because *Cyclospora* infections are frequently heterozygous or multi-strain mixtures that break discrete allele calls, Barratt et al. (2019) bypassed allele catalogues, quantifying pairwise genetic dissimilarity between specimen haplotype sets using an ensemble of Bayesian and heuristic classifiers. This distance grouped 648 of 927 clinical specimens during the 2018 outbreak (Nascimento et al. 2020), matured into a production pipeline with 90% sensitivity and 99% specificity in 2019 (Barratt et al. 2021), and introduced tree-cutting heuristics for cluster detection (Barratt and Plucinski 2023, Jacobson et al. 2023).

While distance-based heuristics established the empirical viability of catalogue-free parasite typing, translating these methods into accessible surveillance tools has proven difficult. The original implementation was developed as an institutional prototype rather than a distributable software package. Consequently, external public health laboratories seeking to adopt the approach had to navigate script releases bound to legacy environments without containerized workflows or test suites (Yanta et al. 2022). Furthermore, because the underlying algorithms were tailored to a specific amplicon panel and pathogen, extending the methodology to other eukaryotic parasites or sequencing assays required bioinformatic re-engineering. To address these limitations, we developed PyEuk to refine the statistical foundations and computational performance of haplotype-based typing. By replacing heuristic classifier ensembles with dropout-tolerant identity-by-state metrics and vectorizing pairwise calculations, PyEuk executes substantially faster while establishing a principled framework for cluster discovery.

Existing amplicon pipelines cleave into two distinct camps. General community-profiling suites such as mothur and QIIME 2 (Schloss et al. 2009, Bolyen et al. 2019) identify whole-amplicon sequence variants without reference coordinate placement, rendering haplotype definitions dependent on read length and primer trimming. Conversely, specialized parasite-typing tools— including SeekDeep, HaplotypR, AmpSeqR, MAD^4^HatTeR, and vivaxGEN—are tailored to specific pathogens or multiplex panels, requiring user-supplied primer coordinates, custom target databases, or organism-specific heuristics (Lerch et al. 2017, Hathaway et al. 2018, Han et al. 2023, Aranda-Díaz et al. 2025, Kleinecke et al. 2025). Similarly, veterinary metabarcoding platforms like Nemabiome depend on curated marker databases for species assignment (Avramenko et al. 2015, Workentine et al. 2020). Adapting existing pipelines to emerging parasites or new panels therefore requires substantial manual effort, often without validated reference schemes (Jesudoss Chelladurai et al. 2025).

To decouple multilocus typing from curated allele catalogues and assay-specific scripts, we developed PyEuk—a modular, open-source software framework and Galaxy workflow ecosystem (Galaxy Community 2024). PyEuk defines and names microhaplotypes deterministically from sequence differences relative to a reference within data-adaptive coverage windows, generating portable labels without centralized databases. When amplicon coordinates are uncharacterized or unavailable, PyEuk reconstructs the panel *de novo* from coverage peaks of raw sequencing reads mapped to a draft genome. Furthermore, PyEuk quantifies clustering uncertainty by evaluating stability across bootstrap resamples, reporting supported cluster count ranges alongside stable transmission cores. We demonstrate PyEuk across five surveillance and clinical validation cohorts from *Cyclospora cayetanensis* and *Plasmodium vivax*, evaluating foodborne outbreak investigations with curated and *de novo* panels, an expanded multi-cluster outbreak assay, a six-year national surveillance archive, a clinical recurrence study, and travel-associated malaria surveillance with panel sizes ranging from 6 to 495 amplicons.

## 2 Results

### An end-to-end framework for catalogue-free typing and panel inference

PyEuk operationalizes multilocus typing through an end-to-end analytical framework implemented as modular command-line tools within the `pyeuk` suite and interoperable Galaxy workflows (Figure 1). Depending on whether amplicon target coordinates are pre-defined or inferred *de novo*, analyses follow one of two pathways. In the Standard workflow, aligned reads are first processed by `define-windows` to delineate shared, high-coverage coordinate intervals across the cohort, ensuring that all specimens are evaluated over an identical reference baseline. Within each window, `call-haplotypes` extracts spanning reads, resolves insertions and deletions, and filters sequencing errors based on read counts and abundance. `PyEuk` avoids arbitrary integer allele numbers: each haplotype receives a deterministic label recording its exact variant positions relative to the reference (for example, = for a reference match, or notation such as `14A>G` or `28_29insT`), allowing laboratories to compare calls directly without synchronizing a central database. These calls are compiled by `build-sheet` into a specimen-by-haplotype matrix that explicitly distinguishes biological absence from missing calls caused by PCR dropout. Finally, `eukaryotyping` computes pairwise heterozygosity-weighted identity-by-state (wIBS) distances across mutually amplified loci, and `cluster` performs hierarchical grouping.

**Figure 1.**
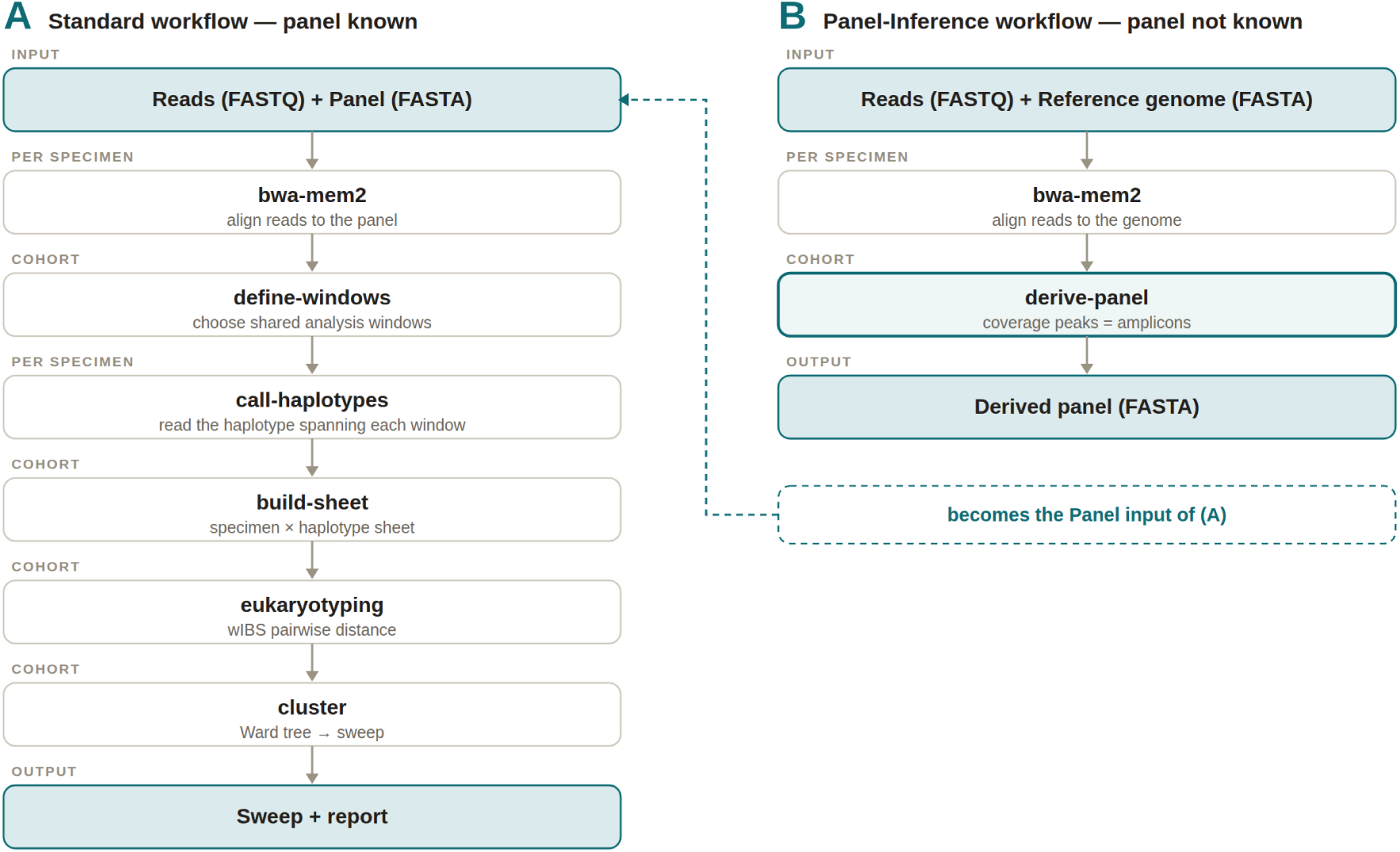
The two PyEuk workflows. **(A) Standard workflow**, used when the panel is known. `bwa-mem2` (Vasimuddin et al. 2019) aligns reads to the panel, and `define-windows, call-haplotypes, build-sheet, eukaryotyping`, and `cluster` produce the sweep and report. **(B) Panel-Inference workflow**, used when the panel is unavailable. `bwa-mem2` (Vasimuddin et al. 2019) aligns reads to the reference genome, and `derive-panel` identifies amplicon coverage peaks and writes their reference sequences to a panel FASTA. This becomes the panel input for (A). The tag above each step indicates whether it operates on one specimen or the full cohort.

Secondary analysis of amplicon archives is routinely hamstrung when primer schemes and target coordinates are omitted from sequence repositories. PyEuk overcomes this through its Panel-Inference workflow, reconstructing target panels *de novo* from raw read coverage. Because targeted amplicon sequencing concentrates reads into sharp depth peaks against an unamplified genomic background, mapping a sample of reads to a draft reference assembly delineates amplified targets. The `derive-panel` module discovers these contiguous peaks via a data-adaptive coverage threshold and exports the corresponding reference sequences into a panel FASTA for downstream analysis. Inferred panels frequently achieve higher and more uniform cohort coverage than the original curated markers (Figure S1).

Hierarchical clustering trees are often cut at fixed thresholds, which can impose artificial boundaries on continuous genetic variation. PyEuk instead evaluates partition stability across bootstrap resamples. Because missing data from PCR dropouts can distort genetic distances, the `cluster` module regularizes pairwise dissimilarities into a valid metric space before constructing a hierarchical tree. When independent clustering metrics concur, the pipeline reports a single partition. When support is divided, it reports the range of supported cluster counts and identifies stable cores—groups of specimens that consistently cluster together across resamples—alongside the fraction of specimen pairs decisively classified as linked or unlinked.

### Resolving foodborne outbreak clusters with curated and de novo amplicon panels

The 2018 multi-state *Cyclospora* outbreak provides a benchmark to test whether PyEuk can resolve real-world transmission chains with and without curated panel coordinates. During this outbreak, epidemiologic traceback linked 153 clinical specimens to two independent commercial produce distributors: 98 cases to Vendor A and 55 to Vendor B (Nascimento et al. 2020). We typed these 153 specimens under two parallel regimes: first using the authors’ curated eight-marker amplicon panel, and second using a panel inferred *de novo* from raw sequencing reads mapped to a draft reference genome. While the curated assay targets eight loci, the eighth marker is a mitochondrial junction with variable-number tandem repeats that requires multi-template matching against twenty junction sequences rather than a single linear reference (Methods §4b). Because PyEuk aligns reads against linear coordinates without variant catalogues, it excludes this repeat junction, typing the remaining seven single-reference markers across 24 sub-amplicon windows.

Applied to the 153 clinical specimens using the curated eight-marker panel, PyEuk’s consensus sweep converges on *k* = 2, separating the two commercial distributors. This partition closely matches the epidemiologic traceback (adjusted Rand index 0.9737), identifying 14 stable cores with 90% of specimen pairs decisively classified as linked or unlinked across bootstrap resamples. All 98 Vendor A specimens cluster into an exclusive group, and 54 of the 55 Vendor B specimens form the second, with one Vendor B specimen assigned to Vendor A. Whereas established surveillance frameworks partition hierarchical trees using supervised distance thresholds tuned against known epidemiologic links (Nascimento et al. 2020; Barratt and Plucinski 2023), PyEuk’s unsupervised sweep recovers this outbreak division directly from sequence dissimilarities without label training.

PyEuk matches this discriminatory accuracy when operating without curated panel coordinates. Mapping raw reads to the draft assembly CcayRef3 (RefSeq GCF_002999335.1; 738 contigs, contig N50 193 kb), `derive-panel` extracted six contiguous coverage peaks. Whereas fixed curated markers exhibit variable amplification efficiency (median depth 588×; Figure S1), `derive-panel` isolates regions of high, uniform coverage (median depth 5,464×), excluding dropout-prone loci. The derived panel separates the two commercial suppliers completely: the primary bifurcation yields an exact 98-and-55 split matching Vendor A and Vendor B (adjusted Rand index 1.0000). Beneath this primary division, higher polymorphism within the derived panel splits the vendor clusters into subclusters; because these internal divisions vary across bootstrap resamples, the sweep reports a range of [12, 28] groups rather than a single point estimate, delineating 15 stable cores with 93% of specimen pairs decisively classified as linked or unlinked.

Reconstructing the typing panel *de novo* preserves the branching structure of the outbreak (Figure 2). When comparing the tree built from the curated eight-marker panel against the tree inferred from the derived six-amplicon panel, pairwise tree distances correlate strongly (cophenetic correlation *r* = 0.9755), and the hierarchical order of cluster merges is preserved (Baker’s Gamma *γ* = 0.9269). Both panels place the same specimens together across all levels of the tree, demonstrating that automated panel discovery captures genuine epidemiological relationships rather than mapping noise. Given only raw amplicon reads and a draft reference genome, PyEuk reconstructs the assay *de novo* and recovers outbreak transmission networks with high fidelity.

**Figure 2.**
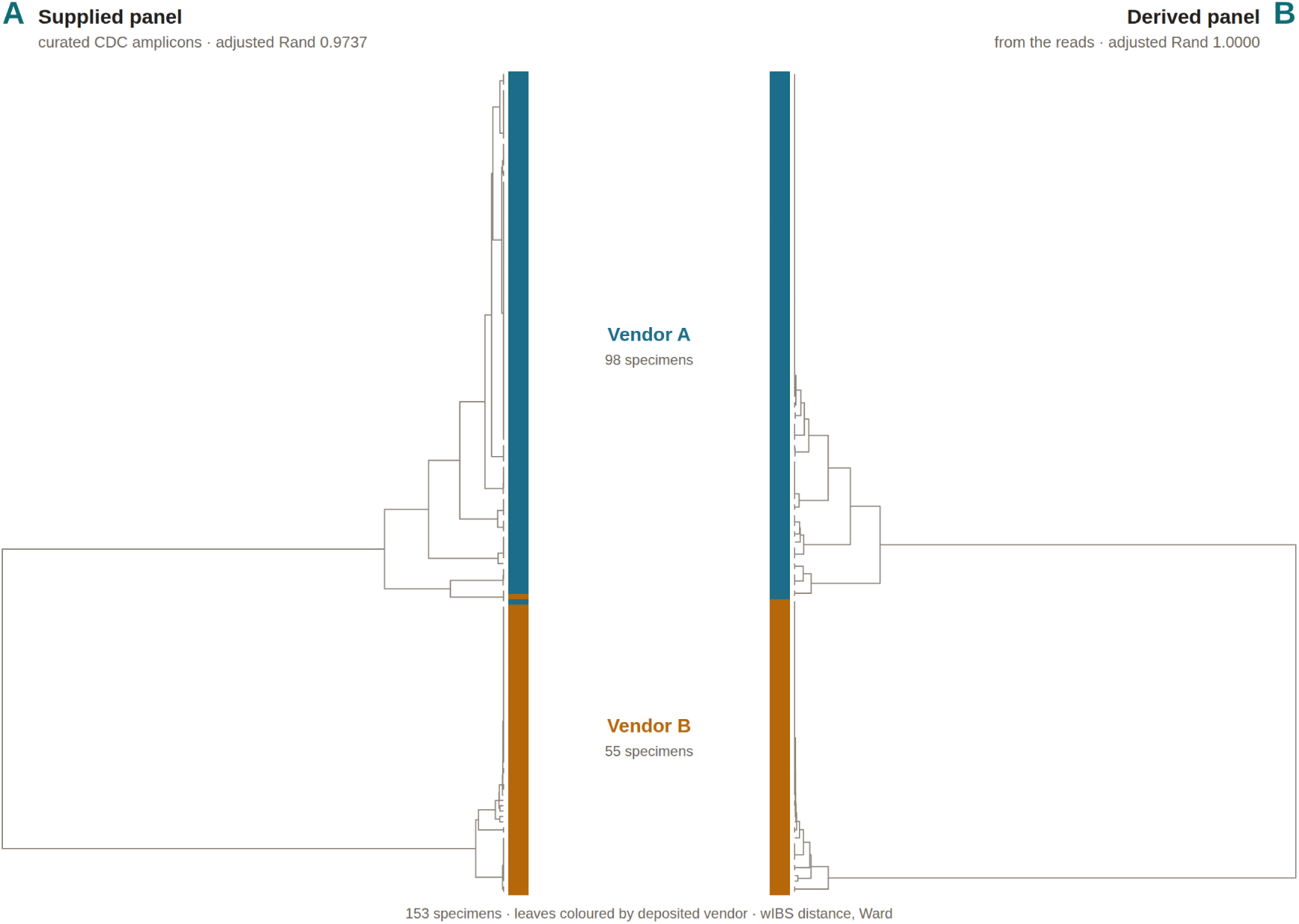
Outbreak cluster recovery and topological congruence between curated and derived amplicon panels. Facing Ward hierarchical trees constructed from pairwise heterozygosity-weighted identity-by-state (wIBS) distances for 153 clinical *Cyclospora cayetanensis* specimens from the 2018 foodborne outbreak. Leaves are coloured by epidemiologically verified tracebacks (Vendor A in blue, Vendor B in orange). **(A) Curated panel**: Using the published eight-marker CDC panel, the primary bifurcation separates the two suppliers with an adjusted Rand index of 0.9737, misplacing only a single Vendor B specimen into the Vendor A cluster. **(B) Derived panel**: Using a six-amplicon panel reconstructed *de novo* by `derive-panel` from raw reads mapped to the draft assembly CcayRef3 without target metadata, the primary bifurcation achieves complete vendor separation (adjusted Rand index 1.0000; 98 Vendor A and 55 Vendor B specimens). Across the full cohort, branching topologies are highly congruent (cophenetic correlation *r* = 0.9755, Baker’s Gamma *γ* = 0.9269), demonstrating that uncurated sequencing reads and a draft reference genome are sufficient to reconstruct transmission chains with high fidelity.

Whereas the 2018 benchmark presented a binary traceback (*k* = 2), molecular surveillance routinely confronts multi-source contamination events where multiple transmission chains circulate concurrently. An independent investigation by the US Food and Drug Administration (Leonard et al. 2024; BioProject PRJNA1052691) provides a benchmark for this regime. Seeking to improve the resolution of the original eight-marker assay, the authors developed an expanded 52-locus targeted amplicon sequencing (TAS) panel and used the Barratt eukaryotyping ensemble to delineate 24 discrete clusters across 66 clinical specimens (CDC01–CDC66). We tested whether PyEuk could recover these clusters without access to the authors’ panel coordinates. Using `derive-panel`, we mapped raw reads to the draft *C. cayetanensis* assembly to reconstruct a 45-amplicon panel *de novo*.

Rather than imposing an arbitrary cut on this multi-cluster population, PyEuk’s unsupervised bootstrap sweep reports a range of [16, 25] groups, bracketing the published count of 24. This stability range holds regardless of whether the analysis is restricted to the focal outbreak cohort or expanded across the entire surveillance repository. Each sequencing library represents an independent patient isolate. Of the 66 clinical specimens evaluated in the published study, 65 pass completeness filtering, yielding 16 stable cores with 95% of pairs decisively resolved. When expanded to all 99 patient libraries in the BioProject archive (84 passing filtering), the sweep yields a consistent structure of 18 stable cores (96% of pairs resolved) across the same [16, 25] range. When evaluated at the published benchmark of *k* = 24, the resulting Ward tree displays strong concordance with the FDA grouping (adjusted Rand index 0.84; Figure 3). Although the two analyses disagree on the broader branching order connecting these clusters (Baker’s Gamma *γ* = 0.63)—as expected when comparing heterozygosity-weighted identity-by-state (wIBS) distances against a classifier ensemble across differing marker sets (45 derived vs. 52 curated loci)—the cluster memberships themselves are preserved. PyEuk thus approximates the multi-cluster architecture reported by the FDA without requiring its curated primer schemes or proprietary classification software.

**Figure 3.**
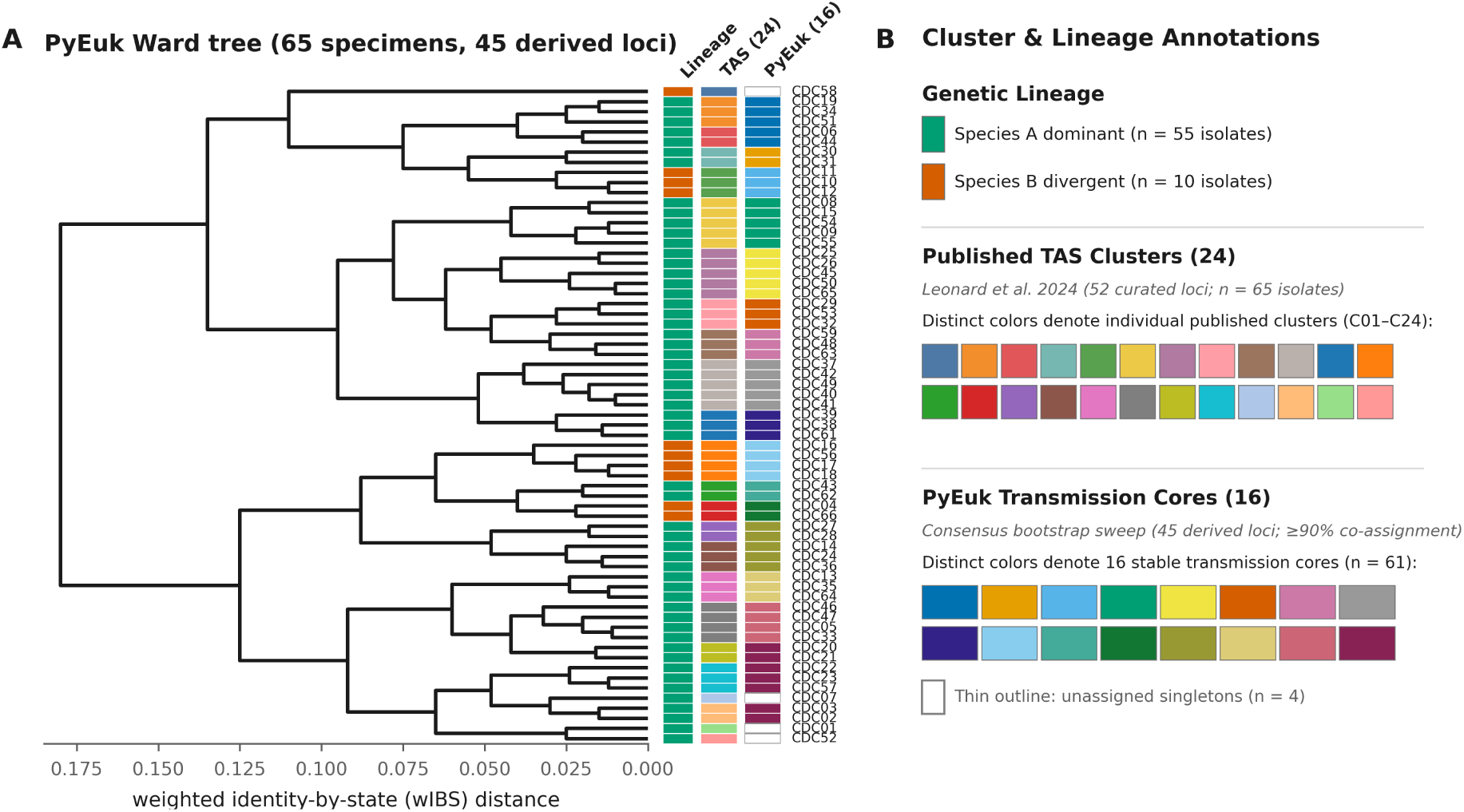
Unsupervised cluster concordance with published multi-cluster outbreak groupings. **(A)** Ward hierarchical tree constructed from pairwise heterozygosity-weighted identity-by-state (wIBS) distances across 65 clinical *Cyclospora cayetanensis* specimens (CDC01– CDC66; PRJNA1052691) using a 45-amplicon panel derived *de novo* without reference panel coordinates. Adjacent color strips indicate genetic lineage (Species A in green, divergent Species B in vermillion), the 24 published targeted amplicon sequencing (TAS) clusters (Leonard et al. 2024), and PyEuk’s 16 stable transmission cores (thin outlines denote singletons outside stable cores). **(B)** Metadata color legend. Genetic lineages denote Species A (*n* = 55, green) and Species B (*n* = 10, vermillion). Swatches denote the 24 published TAS clusters (*C*_01_–*C*_24_; Leonard et al. 2024) and PyEuk’s 16 stable transmission cores (≥ 90% bootstrap co-assignment; thin outlines denote unassigned singletons, *n* = 4). Concordance benchmarks at the published partition (*k* = 24) are adjusted Rand index 0.8408, sweep stability range [16, 25] clusters, 95.2% pairwise resolution, Baker’s Gamma *γ* = 0.6266, and cophenetic correlation *r* = 0.6953.

### Resolving transmission cores across eight thousand national surveillance isolates

Typing frameworks evaluated on small, well-demarcated outbreak clusters often struggle when applied to continuous, large-scale surveillance. To test whether PyEuk can sustain unsupervised cluster inference across continuous public health monitoring, we analyzed the CDC’s six-year national *Cyclospora* genotyping archive (PRJNA578931)—encompassing 8,325 clinical specimens collected between 2018 and 2023 across 40 US states and typed with the original eight-marker assay. This multi-year archive presents an expansive testbed: the data are unlabelled, span multiple transmission seasons, and reflect the biological complexity of sporadic infections, co-circulating outbreaks, and continuous background transmission.

The canonical surveillance assay targets nine loci within our derived 45-amplicon panel. Because each polymorphic locus encompasses multiple data-defined microhaplotype windows, the 8,325 surveillance specimens generated an 8, 325 × 613 window-haplotype matrix—averaging 68.1 distinct haplotypes per locus and capturing circulating allelic diversity alongside multi-strain mixtures. Retaining specimens with haplotype calls across at least 10% of defined windows (min_ completeness ≥ 0.10) excluded 267 low-coverage dropouts, leaving 8,058 high-quality isolates for national-scale clustering. The entire workflow—spanning read mapping, window definition, haplotype calling, distance matrix construction, and a 200-replicate bootstrap sweep—completed in 7.5 hours on a standard 16-core workstation, with the clustering sweep itself processing the 8,058-isolate matrix in approximately 70 minutes (and distance matrix evaluation taking under 4 minutes).

Rather than forcing an arbitrary partition on this multi-year population, PyEuk’s unsupervised bootstrap sweep reports a stability range of [14, 511] clusters—spanning fourteen broad genetic lineages to 511 fine-scale clusters. In the published surveillance literature, investigators routinely enforce a single integer count (*k* = 10 to *k* = 46) by tuning stringency heuristics against known epidemiologic links. When unsupervised selectors are evaluated on these data, however, they diverge markedly (knee, silhouette, and gap criteria do not converge), prompting prior studies to dismiss unsupervised clustering as unviable. In a continuous national transmission reservoir, the cluster count is underdetermined by sequence data alone. Rather than enforcing an arbitrary point estimate, PyEuk’s sweep brackets the literature range while identifying 283 stable transmission cores (≥ 90% bootstrap co-assignment) that decisively resolve 89% of all isolate pairs (Figure 4).

**Figure 4.**
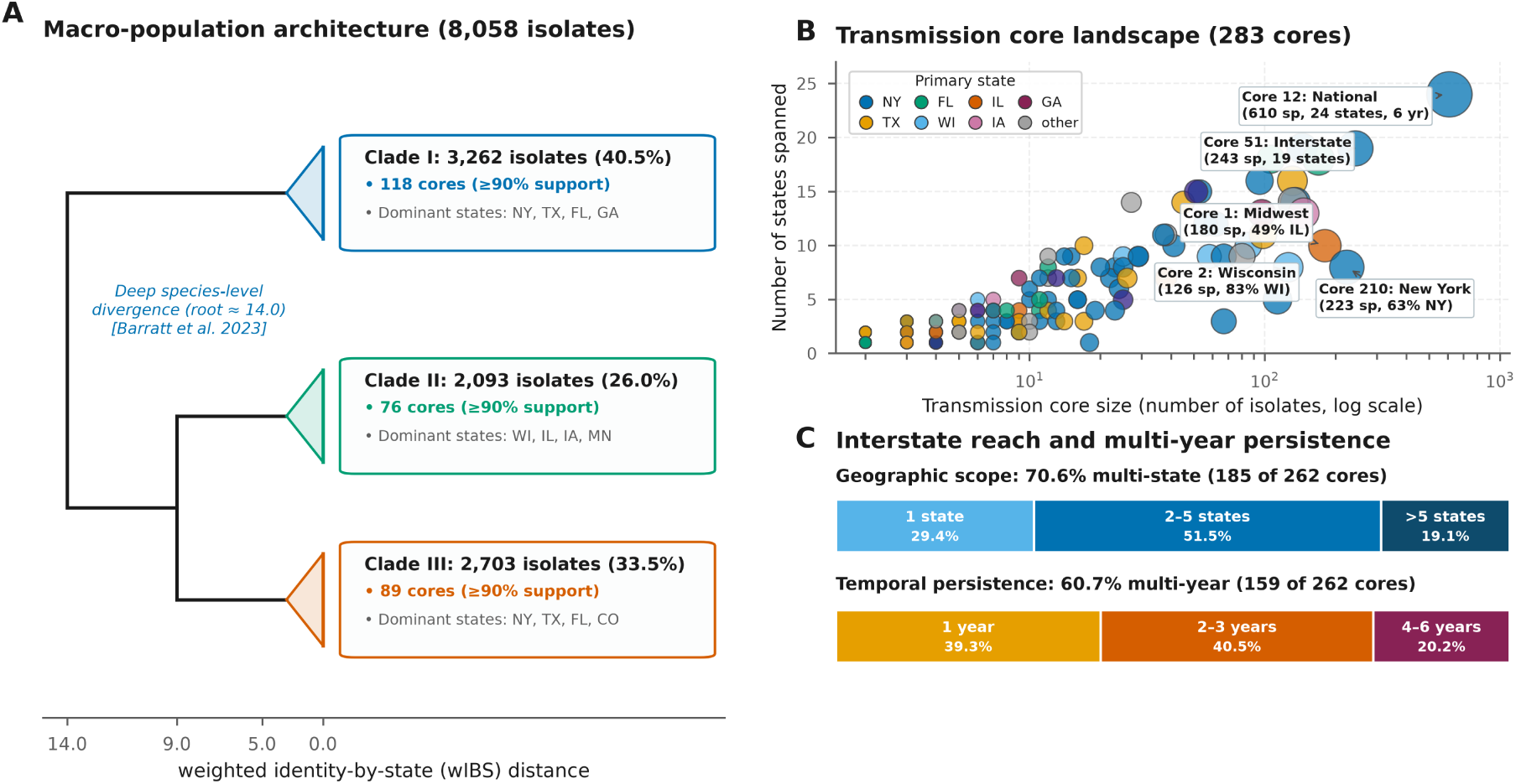
Macro-population structure and transmission core dynamics across eight thousand national surveillance isolates. **(A)** Macro-population architecture across 8,058 clinical *Cyclospora* specimens (PRJNA578931) typed across a derived 45-amplicon panel. Deep divergence at the root (≈ 14.0 wIBS distance) separates the population into three major clades (I–III, corresponding to *Cyclospora* species A, B, and C; Barratt et al. 2023). Within each clade, fine-scale divergence (*<* 2.0) harbors numerous transmission cores (≥ 90% bootstrap co-assignment). **(B)** Transmission core landscape across 283 stable cores, plotting core size (isolate count, log scale) against interstate dispersion (number of US states spanned). Marker size reflects total isolates; colors denote primary reporting state. Callouts highlight landmark transmission archetypes: expansive interstate distribution networks (Core 12 uniting 610 specimens across 24 states and six years; Core 51 spanning 19 states) alongside acute, localized point-source outbreaks (Core 1, 49% Illinois; Core 2, 83% Wisconsin; Core 210, 63% New York). **(C)** Interstate reach and persistence profiles across 262 annotated cores, demonstrating that 70.6% of cores cross state lines (up to 24 states) and 60.7% persist across multiple transmission seasons (2018–2023), directly capturing the dual architecture of national foodborne transmission.

Decoding specimen identifiers confirms that these 283 transmission cores capture epidemiological dynamics rather than technical artifacts. Although public repositories omit case reports, specimen aliases encode reporting state and collection year for 74% of the archive (Methods §13b), annotating 76% of core members across 262 cores. Among these 262 annotated cores, 185 (70.6%) span multiple US states, and 159 (60.7%) span multiple collection years. The largest core encompasses 610 specimens across 24 states over six transmission seasons, consistent with interstate fresh produce distribution networks. Concurrently, PyEuk isolates localized outbreaks (for example, one core with 49% of cases in Illinois, and another with 63% in New York). Unsupervised clustering thus recovers both components of national cyclosporiasis: multi-state distribution networks and localized point-source outbreaks.

### Generalizing multilocus typing across diverse eukaryotic pathogen architectures

Typing frameworks tailored to a single organism frequently encode assumptions—such as uniform diploidy or low within-host diversity—that fail when transferred to divergent eukaryotic architectures. To establish whether PyEuk’s catalogue-free window framework generalizes beyond *Cyclospora*, we evaluated two independent validation cohorts whose biological ground truth was established by external clinical and epidemiological criteria.

The first validation cohort, PvAmpSeq (PRJNA1153071), evaluates *Plasmodium vivax* malaria across an eleven-amplicon microhaplotype panel (*n* = 277). Designed by Rosado et al. (2026), the panel targets eleven of the fourteen *P. vivax* chromosomes (chromosomes 1, 2, 3, 5, 7, 8, 9, 10, 11, 13, and 14), with each locus spanning 89 to 139 base pairs within coding regions of the PvP01 reference genome to capture linked single-molecule microhaplotypes without phasing ambiguity. Here, biological ground truth is defined at two scales. At the macro-geographic scale, truth is established by allopatric continental origin: patient isolates were collected in Peru (*n* = 142) and the Solomon Islands (*n* = 135), representing geographically isolated gene pools with no contemporary exchange. At the micro-epidemiological scale, truth is defined by clinical recurrence adjudication: 91 longitudinally paired patient episodes were independently classified by the clinical trial investigators into homologous relapse (reactivation of dormant liver hypnozoites carrying the identical parasite lineage) versus heterologous reinfection (superinfection from an independent mosquito inoculation). Blind to clinical and geographic metadata, PyEuk recapitulates the continental split (adjusted Rand index 0.9712 against country of origin; 140 of 142 Peru and all 135 Solomon Islands isolates assigned to their continental group) and discriminates relapse from reinfection (ROC AUC 0.9608 across 85 evaluable pairs).

The second validation cohort, CDC AmpliSeq (PRJNA1092573), presents an entirely different design: an open national surveillance archive of 157 travel-associated *P. vivax* cases typed across a high-density 495-amplicon panel. Here, the ground truth is an expert-curated surveillance classification established by CDC investigators using supervised clustering, which resolved the 157 scored specimens into 92 discrete transmission groups—predominantly 79 isolated singletons reflecting sporadic international travel, alongside a modest smattering of small clusters. Without a pre-specified group count, PyEuk’s distance-mode clustering closely reproduces this singleton-dominated benchmark (recovering 93 groups and 83 singletons against 92 deposited groups and 79 singletons). Together, these benchmarks demonstrate that data-defined microhaplotypes sustain robust unsupervised typing across divergent eukaryotic taxa without requiring organism-specific parameter recalibration.

### PvAmpSeq, *Plasmodium vivax*

Controlling *Plasmodium vivax* requires teasing apart whether recurrent malaria is a hypnozoite relapse—reactivation of dormant liver parasites—or a new mosquito reinfection. Unlike *Plasmodium falciparum*, *P. vivax* forms persistent hepatic hypnozoites that can awaken months after successful blood-stage clearance, causing recurrent clinical malaria even in the absence of active vector transmission. The distinction dictates divergent interventions: clinically, a relapse mandates 14-day radical cure with 8-aminoquinolines (such as primaquine or tafenoquine) to clear the cryptic hepatic reservoir; epidemiologically, a reinfection signals ongoing community transmission and the failure of vector control. Because recurrent blood-stage parasites are morphologically indistinguishable by microscopy, microhaplotyping provides an objective basis to adjudicate recurrent episodes.

From the eleven short amplicons developed by Rosado et al. (2026), PyEuk derived 13 catalogue-free windows, generating a 277 by 101 specimen-by-haplotype sheet across 142 Peruvian and 135 Solomon Islands isolates. Operating entirely unsupervised and blind to geographic and clinical annotations, the resulting continuous heterozygosity-weighted identity-by-state (wIBS) metric resolves the parasite population across two distinct biological scales (Figure 5).

**Figure 5.**
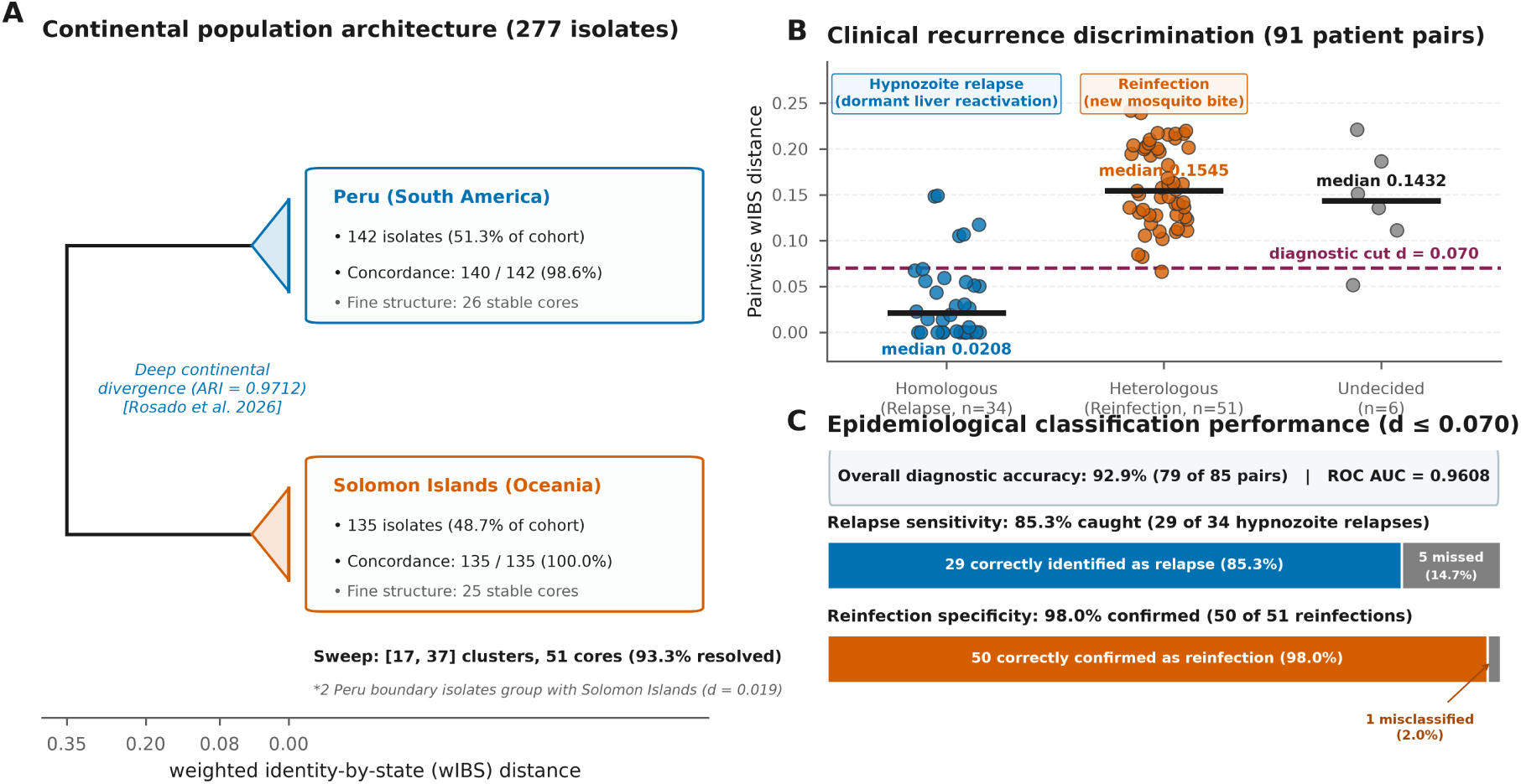
Continental population architecture and clinical recurrence discrimination in *Plasmodium vivax*. **(A)** Continental macro-architecture across 277 clinical *P. vivax* isolates (PRJNA1153071) typed across eleven amplicons. Deep divergence at the root separates the population into two continental branches (ARI = 0.9712 against country of origin), grouping 140 of 142 Peru isolates (blue) and all 135 Solomon Islands isolates (vermillion). Below this primary split, PyEuk resolves 51 stable transmission cores across an unsupervised sweep of [17, 37] clusters (93.3% pairwise resolution). Two boundary Peruvian isolates group adjacent to the Solomon Islands clade (*d* = 0.019). **(B)** Pairwise heterozygosity-weighted identity-by-state (wIBS) distance distributions across 91 longitudinal patient recurrence pairs. Clonal hypnozoite relapses (homologous, *n* = 34) exhibit near-zero divergence (median 0.0208), whereas independent mosquito reinfections (heterologous, *n* = 51) sprawl across the tree (median 0.1545; undecided, *n* = 6, median 0.1432). A diagnostic threshold of *d* ≤ 0.070 (dashed line) cleanly separates reactivation from reinfection. **(C)** Epidemiological classification performance under the diagnostic threshold (*d* ≤ 0.070), achieving 92.9% overall diagnostic accuracy (79 of 85 evaluable pairs, ROC AUC = 0.9608) with 85.3% relapse sensitivity (29 of 34 caught) and 98.0% reinfection specificity (50 of 51 confirmed).

At the macro-geographic scale, PyEuk cleanly separates the two continental populations (Figure 5A). Cutting the dendrogram at two groups segregates 140 of 142 Peruvian isolates into one clade and all 135 Solomon Islands isolates into the other, yielding an adjusted Rand index of 0.9712 against continental origin. The two discordant Peruvian isolates form a tightly linked pair (*d* = 0.019) positioned at the basal boundary between clades. Below this continental split, PyEuk identifies 51 stable transmission cores across an unsupervised stability sweep of [17, 37] clusters, partitioning 93.3% of specimen pairs with high bootstrap support (≥ 90%).

The same genetic distance that separates continents also separates relapses from new infections (Figure 5B). Among the 91 longitudinal patient pairs available in the cohort (out of 106 deposited by Rosado et al. 2026), homologous relapses exhibit near-zero genetic distances (median 0.0208), reflecting clonal reactivation of dormant liver hypnozoites, whereas heterologous reinfections exhibit seven-fold greater divergence (median 0.1545), reflecting independent inoculations from the local parasite pool (ROC AUC = 0.9608 across 85 evaluable pairs; 6 pairs were clinically undecided, median 0.1432).

At an empirical diagnostic threshold of *d* ≤ 0.070, PyEuk achieves 92.9% overall classification accuracy (79 of 85 pairs; Figure 5C), detecting 85.3% of hypnozoite relapses (29 of 34 reactivations) and confirming 98.0% of reinfections (50 of 51 episodes, with one false positive; ROC AUC = 0.9608). Established recurrence classifiers such as the Bayesian framework Pv3Rs or pairwise identity-by-descent models require population allele frequencies and informative priors, and carry false discovery rates of 20% to 27% in sensitivity analyses (Rosado et al. 2026). Operating without external population allele frequency priors or models, PyEuk achieves comparable classification accuracy. Because this threshold was identified on these pairs, prospective cohorts will be needed to fix operational cutoffs. Across eleven short loci spanning 1.3 kb of sequence, PyEuk separates continental lineages and distinguishes relapses from reinfections without organism-specific tuning.

### CDC AmpliSeq, *Plasmodium vivax*

National genomic surveillance of travel-associated malaria presents a distinct epidemiological challenge: distinguishing sporadic international importations from domestic transmission outbreaks. In non-endemic countries like the United States, most malaria diagnoses represent independent travel to diverse endemic regions across the globe. Public health surveillance must therefore solve an asymmetric classification problem: isolating the majority of unrelated singletons while flagging common-source clusters or domestic outbreaks.

The CDC AmpliSeq cohort (PRJNA1092573) tests this capacity at scale. Typed across 495 amplicons distributed across the *P. vivax* genome—more than an order of magnitude larger than the PvAmpSeq panel—the open surveillance archive profiles 157 scored specimens against an expert-curated CDC benchmark. Dominated by sporadic international travel, this benchmark resolves the population into 92 transmission groups, including 79 isolated singletons alongside a modest smattering of small clusters.

This singleton-dominated architecture illustrates the distinction between clustering objectives. Standard hierarchical clustering configured for closed clinical cohorts (count mode) assumes that isolates partition into a small number of cohesive clades; when applied to open surveillance, count sweeps hit artificial group ceilings and merge unrelated travel cases into large clusters. Indeed, PyEuk’s count sweep on the 169 completeness-filtered specimens (derived from 1,960 windows on a 196 by 5,222 matrix) returns [17, 38] clusters with the true count undetermined, correctly signaling that the population cannot be represented by a modest group count.

To match the open surveillance setting, PyEuk switches to distance mode, cutting the hierarchical tree at a fixed dissimilarity threshold (*d* = 0.0869) so that unrelated cases remain singletons. Under this label-free threshold, PyEuk closely mirrors the CDC expert curation without access to external annotations, recovering 93 transmission groups and 83 singletons (adjusted Rand index 0.7801; optimizing the distance cutoff directly against the CDC benchmark yields an upper ceiling of 0.8063; largest group 25 versus 19 deposited). The contrast between PvAmpSeq and CDC AmpliSeq demonstrates that study design dictates clustering strategy: closed clinical cohorts require count mode to resolve shared transmission, whereas open surveillance archives require distance mode to preserve singletons.

Projecting these distance-mode clusters across collection year (2003–2024) and reporting jurisdiction (29 U.S. states) distinguishes potential transmission links from sporadic travel (Figure 6). While most imported cases remain solitary singletons scattered across jurisdictions, PyEuk identifies prominent multi-state transmission lineages. Genotype grp66 connects seventeen specimens diagnosed across seven states during the 2023 season, whereas genotype grp65 links cases spanning five states across eight consecutive years (2017–2024). Because the national archive records jurisdiction of diagnosis rather than country of exposure, unsupervised distance clustering provides a systematic framework to distinguish shared epidemiological exposure from independent travel importations.

**Figure 6.**
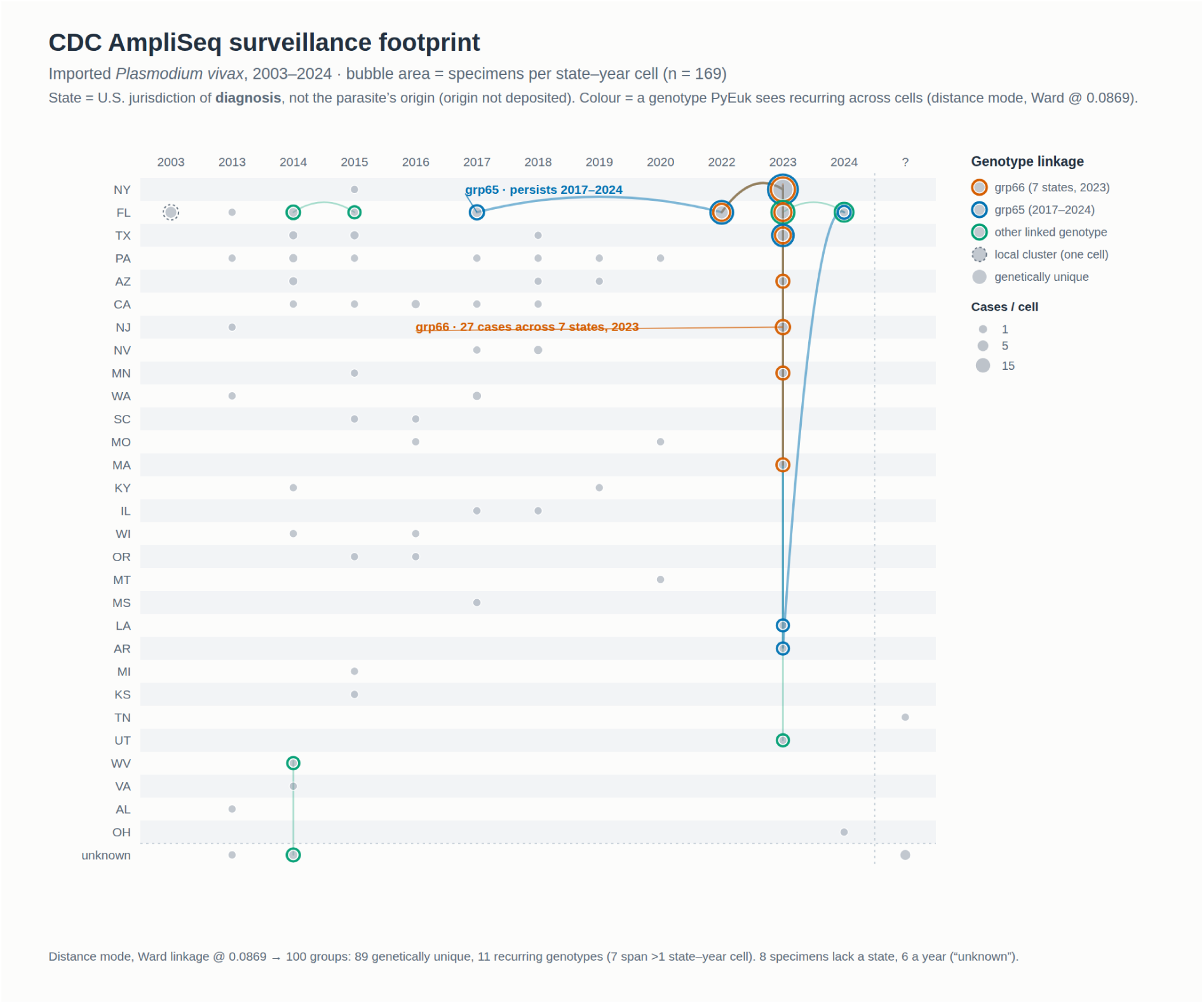
CDC AmpliSeq surveillance footprint. Bubbles show specimens by U.S. state of diagnosis (rows, 29 states) and collection year (columns, 2003–2024). Bubble area represents specimen count (n = 169). State identifies the jurisdiction of diagnosis for an imported case. The archive does not record the parasite’s country of exposure. Rings identify cells containing a genotype assigned to a multi-specimen distance-mode group (Ward linkage cut at 0.0869). **grp66** (vermillion) occurs in seven states in 2023, and **grp65** (blue) occurs from 2017 to 2024 across Florida, Louisiana, Arkansas, New York and Texas. Five further genotypes (green) each link a smaller set of cells. A dashed ring marks a two-specimen cluster within a single cell. Clustering includes all 169 retained specimens (100 groups, including 89 singletons). The adjusted Rand agreement in the text uses the 157 specimens also labelled in the deposited cluster table. Eight specimens lack a recorded state and six lack a year, shown as ‘unknown’.

## 3 Discussion

Molecular surveillance of eukaryotic pathogens has historically adapted frameworks from bacteriology. Bacterial multilocus sequence typing (MLST; Maiden et al. 1998, 2013) established a standard by indexing variation against curated databases of discrete alleles—a system that encounters biological limits in eukaryotic parasites characterized by meiotic recombination, polyclonality, copy-number variation, and within-host diversity. For neglected eukaryotic pathogens, maintaining static allele catalogues places a substantial curation burden on surveillance laboratories. PyEuk addresses this by decoupling microhaplotype extraction from pre-existing catalogues, extracting physically linked variants directly from individual sequencing reads across defined genomic windows. The framework is pathogen- and amplicon-agnostic—scaling from compact surveillance assays in *Cryptosporidium* (Ryan et al. 2021) to 1,960 targets in extensive panels. However, this flexibility operates within biophysical bounds. First, microhaplotypic phasing is anchored to the length of single reads or merged fragments (typically 250 to 500 bp on Illumina instruments), avoiding statistical phasing across unlinked loci. Second, linear window models cannot accommodate non-linear structural rearrangements or variable-number tandem-repeat (VNTR) junctions that vary in copy number and require multi-template matching or fragment sizing. A prominent example is the eighth marker of the CDC *Cyclospora* panel, a mitochondrial repeat junction that cannot be aligned to a single linear reference and is omitted in catalogue-free analysis (Methods §4b). That PyEuk achieves near-perfect epidemiological separation (ARI = 0.9737) using the seven linear loci demonstrates that dense single-molecule microhaplotypes substitute handily for the absent repeat junctions, though integrating specialized repeat callers remains an important future direction. Furthermore, PyEuk operates with or without prior locus coordinates: while it refines user-supplied BED intervals, executing `derive-panel` discovers amplicons *de novo* from raw read coverage peaks, as demonstrated by coordinate-free reconstruction of the *Cyclospora* panel (45 derived of 52 curated loci; Leonard et al. 2024).

Our empirical findings also illuminate broader practical challenges in targeted amplicon panel design for eukaryotic pathogen surveillance. Across multiplex PCR assays, sequencing coverage across targeted ampliconic regions varies wildly (Figure S1). Multiplex primer design for eukaryotic parasites is inherently demanding due to fluctuating GC composition, complex repetitive architectures, and unrecognized polymorphism within primer-binding regions across diverse strains (LaVerriere et al. 2022). Consequently, widely used community panels frequently exhibit suboptimal thermodynamic or stoichiometric balance, resulting in pronounced amplification disparities where a small subset of efficient amplicons sequesters the vast majority of sequencing throughput while other targets achieve only marginal depth or drop out entirely (Figure S1). In the 495-amplicon *P. vivax* CDC AmpliSeq panel, for example, median amplicon depths span several orders of magnitude, with numerous targets receiving only tens of reads. Similarly, in the 2018 *Cyclospora* outbreak cohort, the curated panel yielded a median depth of 588× alongside substantial interquartile dispersion, whereas empirical panel derivation concentrated sequencing effort on consistently amplified coverage peaks (median depth 5,464×; Figure S1). These observations suggest that general amplicon panels are frequently not designed or balanced as effectively as they could be, and would benefit from iterative experimental re-optimization to prune inefficient loci and balance multiplex pools. Until such panel refinements are routine, analytical software must remain robust to these design imperfections. PyEuk accommodates this operational reality both upstream—by enabling data-adaptive panel derivation that selectively captures robustly covered regions—and downstream, by incorporating locus-dropout-tolerant pairwise distance formulations that prevent uneven target performance from biasing phylogenetic and epidemiological inference.

Translating genetic divergence into actionable public health clusters depends on the epidemiological objective. In closed traceback investigations or clinical recurrence trials (such as the 2018 *Cyclospora* outbreak or longitudinal malaria cohorts), cases coalesce into a small, bounded number of exposures; here, count-mode cuts and partition-stability searches identify discrete epidemiological groups. Conversely, in continuous surveillance across wide geographic archives— such as the two-decade CDC AmpliSeq cohort (2003–2024) across 29 U.S. states—the landscape is dominated by sporadic singletons. Rather than forcing a single clustering heuristic across these disparate regimes, PyEuk aligns the tree-partitioning strategy with the study design: count-mode cuts resolve bounded outbreaks, while distance-mode cuts (*d* = 0.0869) preserve background singletons while delineating multi-state transmission clusters. Delineating cluster boundaries in continuous hierarchical trees is an ongoing challenge in molecular epidemiology. The CDC eight-marker *Cyclospora* archive has been analyzed across studies reporting cluster counts from 10 to 46 (Barratt et al. 2021; Barratt and Plucinski 2023; Jacobson et al. 2023; Shen et al. 2025), reflecting the sensitivity of static distance thresholds in an expanding surveillance archive. PyEuk complements these frameworks by reporting stability sweeps rather than prescribing a single cut: with the curated panel, the sweep settles decisively on [2, 2] when boundaries are discrete, while delineating a broader stability range of [14, 511] with 283 reproducible cores across 8,058 national surveillance isolates. Communicating this stability envelope helps epidemiologists distinguish robust transmission cores (≥ 90% bootstrap co-assignment) from boundary-sensitive assignments.

Across cohorts spanning two eukaryotic phyla, PyEuk mirrored or refined published surveillance findings. PyEuk separated the 2018 *Cyclospora* distribution chains (Vendor A 98/98, Vendor B 54/55; adjusted Rand index ARI = 0.9737, 1 error; [2, 2] sweep); mapped 24 published *Cyclospora* TAS clusters without panel coordinates into a stable range of [16, 25] clusters and 16 transmission cores (ARI = 0.8408; Leonard et al. 2024); replicated published CDC AmpliSeq surveillance groupings (93 groups and 83 singletons vs. 92 groups and 79 singletons published; ARI = 0.7801 label-free, 0.8063 tuned); and recapitulated the tripartite *C. cayetanensis* species divergence (*C. cayetanensis* species A, B, and C; Barratt et al. 2023) across 8,058 isolates. In *Plasmodium vivax* (Rosado et al. 2026), PyEuk recovered continental lineages (Peru 140/142, Solomon Islands 135/135; ARI = 0.9712) and classified 85 longitudinal recurrence pairs with 92.9% accuracy (AUC = 0.9608, detecting 29 of 34 relapses with 1 false positive at *d* ≤ 0.070). While Bayesian identity-by-descent algorithms such as Pv3Rs (Rosado et al. 2026) provide formal probabilistic likelihoods, their calibration can be sensitive to background allele frequency estimation in settings with limited population baselines, where false discovery rates of 20% to 27% have been reported. When amplicon panels provide high microhaplotypic diversity, direct observation of physical multi-SNP configurations offers empirical classification without requiring external population allele frequency priors or models, while explicit genealogical models remain valuable for likelihood ratios in low-diversity populations.

Scalable molecular epidemiology requires accessible software architectures that execute rapidly on standard computational infrastructure. PyEuk is an end-to-end Python package (`pyeuk`) providing an upstream panel-inference tool (`derive-panel`) alongside a five-stage core command hierarchy (`define-windows, call-haplotypes, build-sheet, eukaryotyping, cluster`). By isolating haplotype extraction from distance evaluation and vectorizing heterozygosity-weighted pairwise calculations over compressed sparse matrices, the software processes expansive cohorts with dispatch; evaluating pairwise distances and hierarchical clustering across the 8,058-specimen CDC *Cyclospora* archive requires under 4 minutes on a standard 16-core workstation. To support public health laboratories operating with limited computational resources, PyEuk is open source and can be downloaded right now from GitHub (https://github.com/spond/pyeuk) and Bioconda (https://bioconda.github.io/recipes/pyeuk/README.html), containerized in Docker and Singularity formats (ghcr.io/spond/pyeuk), with companion Galaxy workflows (Galaxy Community 2024; Bray et al. 2023) currently under review and to become available from the Intergalactic Workflow Commission (IWC; https://iwc.galaxyproject.org/).

These principles generalize across neglected tropical diseases and veterinary parasitology. In waterborne enteric protozoa like *Cryptosporidium parvum* and *C. hominis*, surveillance has traditionally relied on Sanger sequencing of the hypervariable *gp*60 locus, which provides broad typing but cannot disentangle mixed-strain oocyst co-infections (Ryan et al. 2021), while *Giardia duodenalis* displays allelic sequence heterozygosity and deep assemblage divergence (Cacciò et al. 2008) that challenge standard MLST. Similarly, kinetoplastid parasites like *Leishmania* and *Trypanosoma cruzi* display aneuploidy, somy variation, and hybrid lineages (Rogers et al. 2011; Zingales 2018); because PyEuk avoids forcing read counts into diploid genotypes and evaluates continuous heterozygosity-weighted identity-by-state across presence/absence haplotype matrices, it natively accommodates gene dosage variation and polyclonality. In veterinary parasitology, amplicon deep sequencing of gastrointestinal nematodes tracks benzimidazole resistance mutations in isotype-1 *β*-tubulin (F167Y, E198A/L, F200Y; Avramenko et al. 2015; Workentine et al. 2020; Jesudoss Chelladurai et al. 2025). By requiring reads to span entire windows, PyEuk preserves physical phase across adjacent codons, clarifying whether resistance mutations reside on shared or distinct haplotypes—a capability equally relevant for tracking linked antifungal resistance in emerging fungal pathogens like *Candida auris* (*ERG11*, *FKS1*; Chow et al. 2018).

Targeted amplicon sequencing is increasingly accessible to public health and disease surveillance programs. By grounding inference in physically observed sequencing reads, decoupling analysis from static reference catalogues, and reporting clustering stability across study designs, PyEuk provides a flexible, modular foundation for molecular epidemiology. As genomic surveillance networks expand, catalogue-free microhaplotype analysis offers a portable framework to translate amplicon sequencing into epidemiological insights across diverse eukaryotic pathogens.

## 4 Methods

### 1. Analysis pipeline and command hierarchy

PyEuk performs unsupervised multilocus microhaplotype genotyping, pairwise genetic distance calculation, and hierarchical clustering directly from amplicon sequencing reads and an unannotated reference panel, operating entirely without pre-curated variant catalogues. Catalogue-dependent typing schemes (such as classical MLST or centralized variant registries) founder when confronting hypervariable eukaryotic pathogens that undergo rapid meiotic recombination, extensive polyclonality, or lack dedicated curation consortia. PyEuk resolves this bottleneck by deriving coordinates, variant definitions, and clustering stability directly from sequencing data through six command-line tools: an upstream panel-inference module (`pyeuk derive-panel`) and a core five-stage analysis pipeline:

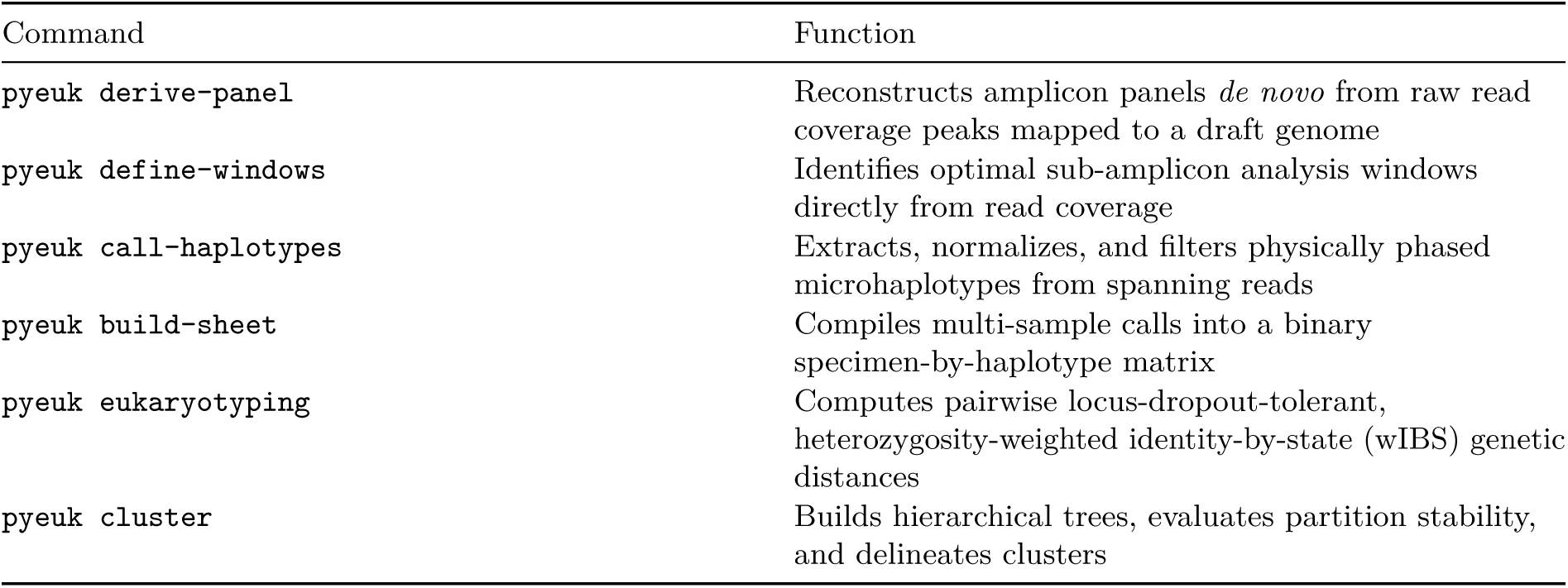

When amplicon coordinates are uncharacterized, `pyeuk derive-panel` extracts candidate markers directly from read coverage; the subsequent three pipeline commands dynamically determine informative sub-amplicon intervals and assemble physical sequence reads into a structured haplotype matrix; the final two commands quantify genetic divergence and partition transmission groups across diverse surveillance study designs (Methods §9 and §10).

### 2. Unit of analysis: linked microhaplotypes versus single-site variants

The fundamental operational unit in PyEuk is the *window microhaplotype*—the exact, gap-normalized nucleotide sequence observed across a defined reference interval on a single, continuous physical sequencing read. Conventional per-site variant callers fragment reads into isolated single-nucleotide polymorphisms, discarding physical phase information and conflating polyclonal mixtures with multiply mutated monoclonal lineages. By strictly conditioning haplotype calls on sequencing reads that cover every nucleotide of a window interval end to end, PyEuk preserves physical within-molecule linkage without invoking statistical imputation or cross-molecule phasing assumptions.

A read contributes to a haplotype call if and only if its alignment spans every base of the target window [*pos*_start_*, pos*_end_]. Across polyclonal infections, this physical constraint preserves true lineage architecture. For example, when two polymorphic sites occur within a single window, spanning reads directly establish whether mutations reside on the same sequenced molecule or represent co-infecting lineages carrying distinct single variants. In a specimen with two variant positions each exhibiting ∼50% allele frequency, a per-site caller cannot distinguish between two co-circulating single-mutant strains and a dual-mutant lineage co-existing with wild type. By conditioning haplotype calls on physical reads that span the entire window end to end (Figure S2), PyEuk directly observes within-molecule phase, distinguishing separate co-infecting lineages from multi-mutant haplotypes without relying on statistical imputation.

### 3. Surveillance and clinical validation cohorts

We evaluated PyEuk across five surveillance and clinical validation cohorts spanning two eukaryotic pathogen species (*Cyclospora cayetanensis* and *Plasmodium vivax*), panel densities from 6 to 495 amplicons, and diverse epidemiological study designs (Table 1). Empirical validation follows the progression of analytical regimes presented in the Results: (1) resolving a binary foodborne outbreak with curated versus *de novo* panels, (2) recovering multi-cluster outbreak structures without reference panel coordinates, (3) scaling unsupervised transmission core discovery across an expansive multi-year national surveillance archive, (4) adjudicating continental divergence and clinical recurrence (relapse vs. reinfection) in malaria, and (5) dissecting singleton-dominated open national surveillance.

**Table 1.** Surveillance and clinical validation cohorts.

| Cohort | BioProject | Panel | Loci | Specimens | Source |
| --- | --- | --- | --- | --- | --- |
| *Cyclospora* 2018 outbreak | PRJNA578931 | CDC 8-marker (7 linear loci used) / Derived 6-amplicon | 7 / 6 | 153 | Nascimento et al. 2020 |
| *Cyclospora* FDA TAS | PRJNA1052691 | Derived 45-amplicon (52-locus TAS) | 45 | 66 (99 archive) | Leonard et al. 2024 |
| *Cyclospora* surveillance | PRJNA578931 | Derived 45-amplicon (9 expanded CDC loci) | 9 | 8,325 (8,058 retained) | Shen et al. 2025;<br>Barratt et al. 2023 |
| *P. vivax* PvAmpSeq | PRJNA1153071 | 11 microhaplotype markers | 11 | 277 | Rosado et al. 2026 |
| *P. vivax* CDC<br>AmpliSeq | PRJNA1092573 | 495 amplicons | 444<br>am-<br>pli-<br>cons<br>(1,760<br>win-<br>dows) | 196 (169<br>retained) | Lancet Reg Health Am 2025 |

Panel sizes range from 6 to 495 sequences across two species, evaluating five distinct surveillance and clinical cohorts:

#### Cyclospora 2018 foodborne outbreak

The 153-specimen 2018 multi-state outbreak cohort represents clinical isolates traced by epidemiologic investigation to two commercial produce distributors: 98 cases to Vendor A and 55 to Vendor B (Nascimento et al. 2020; subset of BioProject PRJNA578931 / PRJNA498911). We typed these specimens under two parallel regimes: first using the published curated eight-marker amplicon panel (typing the seven single-reference linear loci across 24 sub-amplicon windows while omitting the non-linear mitochondrial repeat junction; Methods §4b), and second using a six-amplicon panel inferred *de novo* by `derive-panel` from raw reads mapped to the draft assembly CcayRef3 (RefSeq GCF_002999335.1). This benchmark evaluates binary outbreak cluster recovery (*k* = 2) and topological congruence between curated and derived panels.

#### Cyclospora FDA TAS multicluster cohort

To test multi-cluster resolution in the absence of panel coordinates, we evaluated the targeted amplicon sequencing (TAS) dataset generated by the US Food and Drug Administration (Leonard et al. 2024; BioProject PRJNA1052691). The study established an expert-curated benchmark of 24 discrete clusters across 66 clinical specimens (CDC01–CDC66) using the Barratt eukaryotyping classifier ensemble on a 52-locus panel. We tested whether PyEuk could reconstruct the panel and recover these clusters without access to primer schemes or target coordinates, utilizing `derive-panel` to infer a 45-amplicon panel *de novo* from raw reads mapped to CcayRef3 across the 66 clinical specimens (65 passing completeness filtering) and the full 99-library archive (84 passing filtering).

#### Cyclospora national surveillance archive

To test whether PyEuk sustains unsupervised cluster inference across continuous, large-scale public health monitoring, we analyzed the CDC’s six-year national *Cyclospora* genotyping archive (PRJNA578931; Shen et al. 2025; Barratt et al. 2023). Encompassing 8,325 clinical specimens collected between 2018 and 2023 across 40 US states, this multi-year archive presents an unlabelled, open surveillance testbed spanning multiple transmission seasons. While the 2018 outbreak investigation utilized the original eight-marker assay (seven linear loci evaluated), ongoing national CDC surveillance expanded the scheme to target nine loci within our derived 45-amplicon panel. Filtering at `min_completeness` ≥ 0.10 retained 8,058 high-quality isolates across 613 microhaplotype windows, evaluating macro-population species divergence, bootstrap stability sweeps, and transmission core persistence across states and years.

#### PvAmpSeq, *Plasmodium vivax*

Designed by Rosado et al. (2026; BioProject PRJNA1153071) to investigate recurrent malaria, the PvAmpSeq panel evaluates *P. vivax* malaria across eleven microhaplotype amplicons targeting eleven chromosomes of the PvP01 reference genome (generating 13 catalogue-free windows). The cohort comprises 277 clinical isolates evaluated against two independent ground truths: (1) macro-geographic allopatric origin between Peru (*n* = 142) and the Solomon Islands (*n* = 135), and (2) clinical recurrence adjudication across 91 longitudinally paired patient episodes classified by trial investigators into homologous hypnozoite relapses (*n* = 34), heterologous mosquito reinfections (*n* = 51), and undecided cases (*n* = 6). This cohort tests whether continuous genetic distances can separate continental lineages and distinguish relapse from reinfection without external population allele frequency priors or model tuning.

#### CDC AmpliSeq, *Plasmodium vivax*

The CDC AmpliSeq cohort (PRJNA1092573; Lancet Reg Health Am 2025) evaluates open national genomic surveillance of travel-associated *P. vivax* malaria. Comprising 196 patient libraries collected between 2003 and 2024 across 29 US states and typed across a high-density 495-amplicon panel (444 amplicons passing quality filters and yielding 1,760 retained sub-amplicon windows out of 1,960 published BED intervals), the dataset represents an asymmetric classification benchmark. Retaining 169 specimens with completeness ≥ 0.70, isolates were evaluated against an expert-curated CDC benchmark resolving 157 scored specimens into 92 transmission groups (dominated by 79 isolated singletons). This cohort tests PyEuk’s distance-mode clustering for isolating sporadic international travel importations while identifying multi-state domestic transmission clusters.

All raw sequencing reads were retrieved from the European Nucleotide Archive (ENA). We retained only sequencing runs where both paired-end FASTQ files downloaded completely, verified by comparing downloaded byte counts against archive manifests. Each result set details excluded runs. Amplicon and read lengths are omitted because they could not be independently sourced from public archive metadata.

### 4a. Panel coordinate curation and linear verification

Reference panels were assembled as FASTA files containing one representative sequence per amplicon locus, derived from the source publications’ deposited data, and subjected to coordinate verification prior to alignment. Unsupervised microhaplotyping requires unambiguous coordinates; panel curation resolves indexing discrepancies, identifies non-linear repeat junctions, and screens read archives before alignment.

For PvAmpSeq, cross-referencing the published Chr09 marker (reported as 128 bases in the publication’s table) against the PvP01 reference genome interval (`PvP01_09_v1:1459600-1459728`, 129 bases) identified a missing guanine at zero-based offset 119. All 312 published Chr09 haplotypes are 129 bases long and retain this guanine, demonstrating that the error resided in the publication’s summary table rather than the underlying dataset. We corrected the panel sequence accordingly and retained the original file.

### 4b. Non-linear tandem-repeat junctions

Markers characterized by non-linear repetitive junctions represent an intrinsic methodological boundary for linear catalogue-free microhaplotyping. Standard sequence alignment and window definition assume collinear mapping against a single static reference sequence. When an assay targets variable-number tandem-repeat (VNTR) arrays or complex genomic junctions that vary in repeat copy number, forcing reads against a single linear reference generates clipped alignments, degenerate gaps, and spurious variant calls. Such loci historically demand multi-template sequence catalogues or fragment-sizing algorithms that lie outside the scope of window-based microhaplotyping.

The CDC eight-marker *Cyclospora* panel provides a clear illustration of this architectural constraint: its eighth marker spans a mitochondrial junction containing variable tandem repeats of a 15-base unit, which the original surveillance pipeline types by length classification and multi-template matching against 20 curated junction sequences rather than alignment to a single reference (Nascimento et al. 2020; Barratt et al. 2021). Because PyEuk models linear alignments against a single reference per locus, this marker was omitted, restricting analysis to the seven single-reference loci. This restriction matches the published CDC BED schema, which tiles 24 target regions across these exact seven loci. As demonstrated in the Results, dense microhaplotypes across the seven linear markers provide ample phylogenetic signal to achieve near-perfect outbreak separation (ARI = 0.9737) without the repeat junction.

### 5. Read alignment and primary filtering

PyEuk aligns raw paired-end sequencing reads against the amplicon panel and extracts high-confidence, non-redundant primary alignments. Amplicon libraries frequently contain non-specific primer binding, chimeric artifacts, and multi-mapping reads; alignment filtering ensures that each sequenced molecule is counted once, preventing coverage inflation and spurious haplotype calls.

Reads are mapped using `bwa-mem2` (v2.2.1; Vasimuddin et al. 2019). Alignments are filtered via pysam to retain reads satisfying three strict criteria: (1) mapping quality score MAPQ ≥ 20, (2) flag indicating a properly paired read pair (`PROPER_PAIR`), and (3) exclusion of secondary (`SECONDARY`) and supplementary (`SUPPLEMENTARY`) alignments. Restricting the analysis to primary alignments prevents chimeric read pairs and split alignments from contributing redundant haplotype observations.

### 6. Dynamic catalogue-free window definition

The `pyeuk define-windows` command automatically identifies optimal, strand-balanced sub-amplicon coordinate intervals directly from empirical read alignments without requiring pre-annotated locus boundaries. In targeted amplicon sequencing, forward and reverse sequencing reads originate from disparate primer positions and overlap asymmetrically. Naively pinning an analysis window to the first covered nucleotide restricts spanning coverage to a single strand, whereas maximizing width alone selects intervals that starve one read orientation. Dynamic optimization ensures windows capture maximum haplotype length while preserving balanced, high-depth spanning coverage across both sequencing strands.

Window definition evaluates three properties across each reference sequence: coverage extent, placement, and width. The coverage extent spans the contiguous interval [*pos*_first_*, pos*_last_] exhibiting read depth ≥ 20. Within this extent, candidate intervals are scored by the objective function *S* = width × *f*_span_, where *f*_span_ is the fraction of overlapping reads that span the interval end to end (*f*_span_ ≥ 0.30). Spanning fractions are measured directly from empirical CIGAR alignments rather than inferred from nominal read lengths, avoiding percentile-based estimation biases where a nominal 70% read-length rule delivered only 30% to 33% observed spanning coverage.

Asymmetric read initiation across strands demonstrates the necessity of strand-aware placement. Across an amplified target where forward and reverse reads initiate at staggered positions (for example, spanning positions 191 to 515 with depth ≥ 20, where forward reads initiate predominantly at position 195 and reverse reads at 212), only a small fraction of reads reach the first covered base (107 reads). Tiling from the first base limits the window to 150 bases (48.1% spanning coverage). Scoring width alone selects a 300-base window spanned by 34.2% of reads overall but only 0.04% of forward reads. Multiplying width by spanning fraction shifts the start by 24 bases to position 215 (reached by 92.8% of reads), recovering a 270-base window with 97.6% spanning coverage. The algorithm tiles the remaining coverage extent at this width, placing a boundary at the start of the spannable interval; leading or trailing remnants meeting the minimum width threshold are retained as distinct adjacent windows.

### 7. Microhaplotype calling and abundance-based error filtering

The `pyeuk call-haplotypes` command extracts, normalizes, and filters single-molecule micro-haplotypes across all defined windows for each specimen, reporting observed variants alongside explicit uncalled records. PCR amplification and Illumina sequencing introduce base substitutions and indel alignment ambiguities that artificially inflate haplotype diversity if unpoliced. Pre-count normalization prevents redundant indel representation, while abundance-ratio clustering purges PCR chimeras and single-base sequencing errors without sacrificing authentic low-frequency lineages.

For each specimen and window, the caller: (1) selects reads covering every base of the window, (2) left-aligns all gap coordinates to their leftmost equivalent positions, (3) tallies exact sequence occurrences, (4) purges putative single-base sequencing errors if a 1-bp neighbor is ≥ 8-fold more abundant, (5) applies the tripartite depth and frequency thresholds in Table 2, and (6) writes one record per passing haplotype.

**Table 2.** The three filters of the caller.

| Filter | Function | Default |
| --- | --- | --- |
| `min_span` | Reads that span the window, before any haplotype is examined | 30 |
| `min_reads` | Reads that carry one haplotype | 10 |
| `min_freq` | Fraction of the spanning reads that carry one haplotype | 0.05 |

Leftmost gap normalization precedes sequence counting so that alternative alignments of identical insertion or deletion events collapse into a unified variant tally rather than being recorded as distinct alleles. Surviving haplotypes are designated by 1-based reference substitutions (e.g., `45T>A,57T>C`) or = for reference identity. If no haplotype passes filtering, the locus is recorded as `NOT_CALLED` with its spanning read depth (*N*_span_ *<* 30 denotes uncalled; *N*_span_ ≥ 30 denotes evaluated but filtered), preserving failed specimens in downstream matrices rather than silently dropping them.

### 8. Specimen-by-haplotype matrix construction

The pyeuk build-sheet command compiles multi-specimen haplotype calls into a binary specimen-by-haplotype matrix alongside a long-format read frequency table. Genetic distance calculation requires decoupling allelic presence from observational absence, ensuring that technical locus dropout is treated as missing data rather than identity with the reference.

The matrix assigns specimens to rows and distinct window-haplotypes to columns. An entry of X denotes presence; all columns within a window remain empty if the locus was `NOT_CALLED`. Reference-identical alleles are explicitly tracked under =. An accompanying long-format table records absolute read counts and within-host frequencies for all called variants, preserving quantitative mixture proportions. Cohort-wide minor allele frequency filtering (`min_maf`) is disabled by default (0.0), as applying arbitrary thresholds (such as 0.05) removes informative rare outbreak markers, causing 2,443 of 3,392 successfully called *Cyclospora* windows to appear uncalled (yielding 51 columns when disabled vs 15 columns when filtered at 0.05).

### 9. Weighted identity-by-state genetic distance

The `pyeuk eukaryotyping` command calculates pairwise genetic distances between specimens using a locus-dropout-tolerant, heterozygosity-weighted identity-by-state (wIBS) formulation. Unsupervised surveillance data exhibit heterogeneous amplification dropout across loci. Unweighted distance metrics disproportionately reward matches at uninformative, invariant reference alleles, while standard Euclidean calculations cannot accommodate missing loci without distortion.

For specimens *i* and *j*, pairwise distance is computed strictly over the intersection of mutually called windows *W_ij_*:

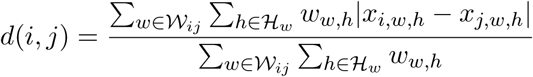

where *x_i,w,h_* ∈ {0, 1} indicates haplotype presence, and weights follow expected heterozygosity:

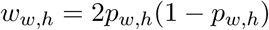

with *p_w,h_* denoting the cohort-wide frequency of haplotype *h* across specimens called at window *w* (the default `heterozygosity` scheme in PyEuk 0.6.0). Ward’s hierarchical clustering assumes Euclidean dissimilarity. Pairwise locus dropout can introduce mild non-Euclidean distortions (negative eigenvalues in the distance Gram matrix). PyEuk provides an optional projection onto the positive semi-definite (PSD) cone by clipping negative eigenvalues to zero. Projection was disabled for count-mode analyses (retaining raw metric geometry) and enabled for high-density distance-mode surveillance (CDC AmpliSeq; Methods §10).

### 10. Hierarchical clustering and partition selection

The pyeuk cluster command performs agglomerative hierarchical clustering under Ward’s minimum variance criterion, interrogates partition stability via a multi-selector sweep, and delineates clusters in either count or distance mode. No single clustering criterion universally fits all epidemiologic paradigms. Closed foodborne outbreaks require partitioning into a discrete number of cohesive transmission groups, whereas national surveillance archives are dominated by unrelated travel singletons where forcing specimens into a low *k* manufactures false epidemiological linkage.

The clustering sweep evaluates candidate cluster counts *k* ∈ [2*, k*_max_] across four independent, label-free criteria: (1) merge-height gap knee (identifying the maximum drop in successive dendrogram join heights, beyond which merging groups adds disproportionate distance), (2) average silhouette width (favoring counts where specimens are closest to their own group relative to the nearest alternative), (3) the gap statistic relative to a uniform null (Tibshirani et al. 2001; comparing within-group spread with that expected in unstructured data), and (4) bootstrap resampling stability (identifying groupings that recur across resamples). Concordance indicates a determined natural partition; divergence flags multi-scale structure, for which PyEuk extracts *stable transmission cores*—maximal subgraphs of specimens co-clustering with ≥ 90% bootstrap support.

Count mode selects an optimal *k* from the largest merge-height gap satisfying two guarded constraints: relative gap Δ*h/h*_max_ ≥ 0.22, and minimum cluster membership *n*_min_ ≥ max(2, min(5, ⌊0.10*n*⌋)). These constants were calibrated on six *Cyclospora* sheets and 12 column-shuffled null cohorts (*k* = 1 for nulls, *k* = 2 for real sheets). Specimens in sub-threshold clusters remain unassigned. Count mode suits closed investigations where every accepted specimen belongs to an exposure group and clusters of two cases are uninformative.

Distance mode cuts the dendrogram at an absolute dissimilarity threshold *d*_cut_ without selecting *k* or searching for a gap, allowing singletons to remain isolated. Without labels, PyEuk sets *d*_cut_ to the empirical 5th percentile of pairwise distances; with labeled training pairs, it sets *d*_cut_ = median(*d*_within_) + 3 × 1.4826 × MAD(*d*_within_).

Operational mode must be selected based on study design, not organism biology. Six label-free distribution tests (Silverman multi-modality, gap statistic, silhouette, nearest-neighbor fractions, median-to-first-percentile ratios, and negative eigenvalue mass) failed to separate CDC surveillance from outbreak cohorts. Across all 157 CDC specimens at baseline completeness (≥ 0.10, including 79 singletons), distance mode achieves an ARI of 0.7952 (rising to 0.8063 when filtering completeness ≥ 0.70, or 0.7801 under label-free cut selection) while count mode collapses to 0.0470 (forcing 79 singletons into one cluster). Conversely, restricting analysis to the four largest published clusters removes singletons, inverting performance: count mode achieves an ARI of 1.0000 while distance mode yields 0.1137.

### 11. Software architecture, containerization, and Galaxy workflows

PyEuk is implemented in Python 3.10+ as an open-source software suite with automated unit tests (54 of 54 passing), containerized images, and Galaxy workflow integrations. Deploying surveillance tools across public health laboratories requires platform independence, containerization, and accessible workflow execution.

Computational bottlenecks in pairwise wIBS distance calculation and dendrogram evaluation are vectorized via NumPy and SciPy. Core read alignment parsing uses `pysam` (packaged via the `amplicon` extra on PyPI). The software is open source and can be downloaded right now from GitHub (https://github.com/spond/pyeuk) and Bioconda (https://bioconda.github.io/recipes/pyeuk/README.html), with container images distributed via GitHub Container Registry (ghcr.io/spond/pyeuk).

Conceptually, PyEuk operates under two overarching analytical workflows (Figure 1): the Standard workflow when amplicon coordinates are known, and the Panel-Inference workflow when panels must be derived *de novo* from raw reads using `derive-panel`. For execution on compute infrastructure, PyEuk’s core Standard pipeline is integrated into Galaxy through three standardized, Planemo-tested workflow implementations: (1) end-to-end FASTQ processing with dynamic window derivation, (2) FASTQ processing with user-supplied BED coordinates, and (3) BAM execution for rapid parameter re-sweeps without repeating alignment. Companion Galaxy workflows are registered on WorkflowHub (Gustafsson et al. 2025), are currently under review, and will become available from the Intergalactic Workflow Commission (IWC; https://iwc.galaxyproject.org/). Each workflow exports a standardized artifact bundle: coordinate BED, specimen-by-haplotype matrix, wIBS distance matrix, cluster membership table, and run-parameter JSON.

### 12. Cohort-specific parameterization and completeness filtering

Detailed parameter configurations applied across empirical validation benchmarks are summarized in Table 3. Panel density and sequencing depth dictate specific quality control thresholds, particularly when distinguishing low-depth technical dropouts from authentic biological variation.

**Table 3.** The settings of each run.

| Cohort | min_span | min_freq | Windows | Projection | Cut mode | min_completeness |
| --- | --- | --- | --- | --- | --- | --- |
| *Cyclospora* | 30 | 0.05 | from reads | off | count | 0.10 |
| PvAmpSeq | 30 | 0.05 | from reads | off | count | 0.10 |
| CDC AmpliSeq | 30 | 0.10 | published BED | on | distance 0.0869 (label-free) / 0.080 (tuned) | 0.70 |

Both *Cyclospora* and PvAmpSeq utilized identical front-end filtering (*N*_span_ ≥ 30, *f*_hap_ ≥ 0.05) and read-derived windows with PSD projection disabled. CDC AmpliSeq applied *f*_hap_ ≥ 0.10 across 1,960 published BED intervals (retaining calls across 1,760 windows; median width 68 bases, range 16 to 78 bases) with PSD projection enabled. The CDC AmpliSeq distance cut of 0.08 maximizes agreement with published surveillance labels; label-free calibration selects 0.0869 and gives an ARI of 0.7801, indicating that 0.08 is label-tuned.

For CDC AmpliSeq alone, `min_completeness` was raised from 0.10 to 0.70. Specimen completeness exhibited a bimodal distribution with an empty gap between 0.5301 and 0.7068 (median 0.945): 169 specimens were at or above 0.70 (minimum 0.7068) and 27 below (maximum 0.5301). Raising the threshold excluded 27 low-coverage specimens (8 with zero calls, 5 below 0.10, 4 between 0.10 and 0.35, and 10 between 0.35 and 0.55), increasing ARI by 0.0111 (from 0.7952 to 0.8063). Completeness filtering is panel-dependent; a cohort covering only part of an assay may exhibit completeness near 0.50 and would be excluded by a 0.70 threshold.

### 13a. Performance evaluation and benchmark scoring

We evaluated clustering accuracy and distance discrimination against independent ground truth benchmarks from source publications (Table 4). Unsupervised typing must be rigorously validated against external clinical, geographic, and outbreak annotations without using those annotations during inference.

**Table 4.** Ground truth benchmarks for empirical scoring.

| Cohort | Comparison | Source |
| --- | --- | --- |
| Cyclospora 2018 outbreak | Two commercial distributor clusters | Epidemiologic traceback (Nascimento et al. 2020) |
| Cyclospora FDA TAS | 24 multi-cluster outbreak groupings | Published classifier ensemble (Leonard et al. 2024) |
| Cyclospora national surveillance | Species clades and multi-state/year core persistence | Archive metadata and inferred state/year (Shen et al. 2025; Barratt et al. 2023) |
| PvAmpSeq | Continental origin (Peru vs. Solomon Islands) | Archive metadata |
| PvAmpSeq | Relapse versus reinfection across 91 recurrence pairs | Clinical trial adjudication (Rosado et al. 2026) |
| CDC AmpliSeq | 92 transmission groups (79 singletons) and travel history | Expert-curated CDC benchmark (Lancet Reg Health Am 2025) |

Partition concordance was evaluated via the Adjusted Rand Index (ARI), where 1.0 indicates identical assignments and 0 indicates chance-level agreement. Each ARI is reported alongside the number of scored specimens, excluding unassigned isolates. Genetic distance discrimination was quantified by Receiver Operating Characteristic Area Under the Curve (ROC AUC), where 0.5 denotes no discrimination and 1.0 denotes complete separation. Comparisons utilize deposited results from source publications without rerunning external tools in frozen histories.

### 13b. Inferring specimen state and year from identifiers

Public sequencing archives frequently omit case report forms and epidemiological metadata. In the absence of clinical records, structured sample identifiers provide an independent means to evaluate whether transmission cores capture authentic multi-state or localized outbreaks.

In BioProject PRJNA578931, sample aliases encode reporting state and collection year within structured naming schemas. Aliases follow `<yy>US<ST>…`: a two-digit year, country code `US`, and two-letter state abbreviation (e.g., `18USIA…` denotes Iowa 2018). Regex parsers extracted state and collection year for 6,136 of 8,325 specimens (74%) in PRJNA578931 (annotating 76% of core members across 262 cores).

These annotations represent inferred metadata rather than formal epidemiological records: reporting state does not guarantee the place of exposure, and collection year cannot resolve discrete transmission chains within a season. Because multi-state produce distribution produces multi-state *Cyclospora* outbreaks, cores spanning multiple states are expected under commercial foodborne transmission. We therefore use state and year to describe the geographic and temporal composition of stable transmission cores, never as ground truth for ARI scoring.

### 14. Held-out cross-validation and parameter stability

We evaluated the generalization performance and parameter stability of the minimum spanning read filter (`min_span`) across held-out test partitions. Because the full-cohort benchmark threshold of `min_span` = 30 was initially identified via a parameter sweep across the 153 *Cyclospora* outbreak specimens, repeated cross-validation is required to quantify potential in-sample optimization bias.

We executed 40 independent random Monte Carlo splits (65% training, 35% testing; *n*_test_ = 53). For each split, we evaluated two settings on the held-out specimens: a fixed default of `min_span` = 30 and a value tuned strictly on that split’s training set. We also recorded the value obtained by tuning on the held-out set itself, representing an empirical upper-bound oracle.

A fixed threshold of 30 yields a mean held-out ARI of 0.8608, a median of 0.9237, and a standard deviation of 0.2594 (with three of the 40 splits returning 0.0000 due to over-pruning). Training-set tuning achieves a mean ARI of 0.8950 and a median of 0.9242. The oracle mean is 0.9524, representing an in-sample selection advantage of 0.0574.

## 15. Data and code availability

All sequencing datasets analyzed in this study are publicly available from the NCBI Sequence Read Archive (SRA) and European Nucleotide Archive (ENA) under the BioProject accessions listed in Table 1. PyEuk is open source and can be downloaded right now from GitHub (https://github.com/spond/pyeuk) and Bioconda (https://bioconda.github.io/recipes/pyeuk/README.html), with container images distributed via GitHub Container Registry (ghcr.io/spond/pyeuk). Companion Galaxy workflows are currently under review and will become available from the Intergalactic Workflow Commission (IWC; https://iwc.galaxyproject.org/).

## 5 Acknowledgments

We thank the entire Galaxy team and Björn Grüning for their support and contributions to this work. This research was supported by the National Institute of Allergy and Infectious Diseases (NIAID) of the National Institutes of Health (NIH) under award number [U24AI183870](https://reporter.nih.gov/search/ITDZqr1H70e4HYCRe8gT6A/project-details/11141082) (*An integrated platform for multiomic analyses of pathogen and host data using scalable public infrastructure*).

## 7 Supplementary figures

**Figure S1.**
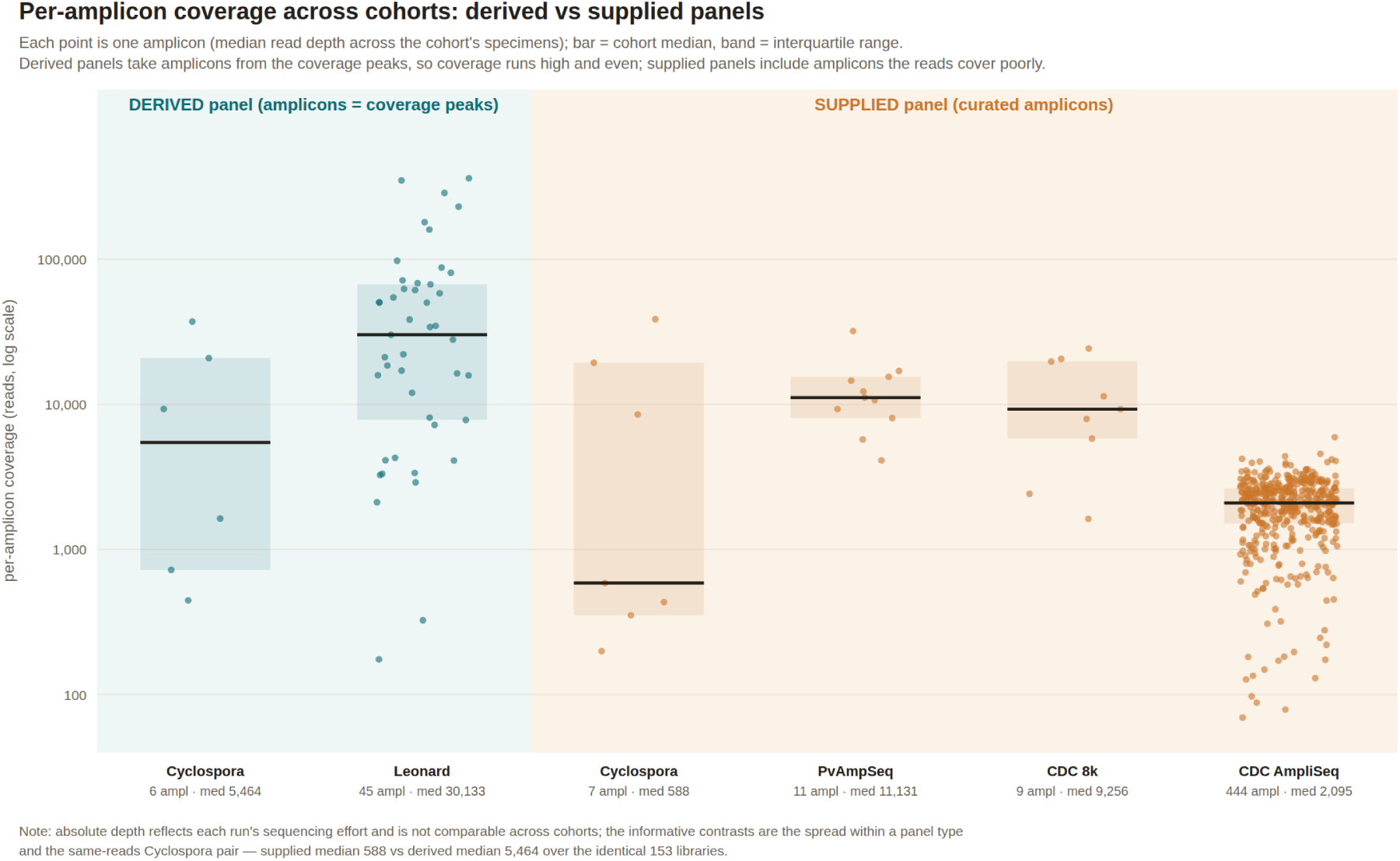
Per-amplicon coverage across cohorts, derived versus supplied panels. Each point shows an amplicon’s median read depth across specimens. Bars mark cohort medians and bands mark interquartile ranges. Depth is plotted on a log scale. Derived panels (left) select amplicons from coverage peaks and have high, even coverage. Supplied panels (right) include poorly covered amplicons and show lower, more variable depth, particularly the 495-amplicon CDC AmpliSeq panel, where some amplicons have only tens of reads. In the clearest comparison, both *Cyclospora* panels use the same 153 libraries: median per-amplicon coverage is 588 for the supplied panel and 5,464 for the derived panel, which selects the covered regions. Absolute depth depends on sequencing effort and is not comparable across cohorts. The relevant comparison is variation within and between panel types.

**Figure S2.**
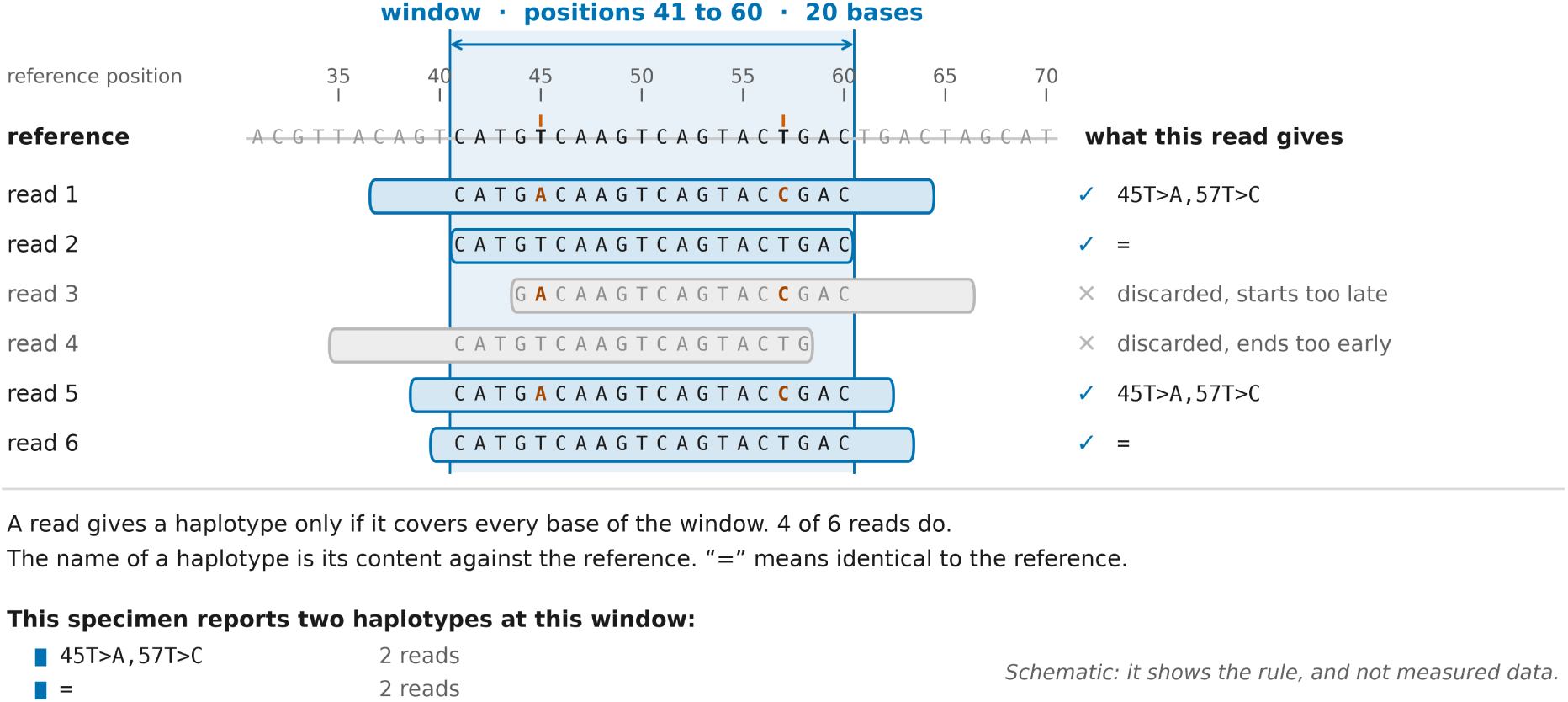
The window haplotype. A window is an interval on a reference sequence. Only reads covering every base contribute a haplotype: read 3 starts inside the window and read 4 ends inside it, so both are excluded. Each of the other four reads contributes one haplotype. Names describe differences from the reference: `45T>A,57T>C` denotes two differences, and = denotes a reference match. The specimen has two haplotypes at this window, each supported by two reads. This is a schematic, not measured data.

## Notes

### Competing Interest Statement

The authors have declared no competing interest.

